# A single spectrum of neuronal identities across thalamus

**DOI:** 10.1101/241315

**Authors:** James W. Phillips, Anton Schulmann, Erina Hara, Chenghao Liu, Lihua Wang, Brenda C. Shields, Wyatt Korff, Andrew L. Lemire, Joshua Dudman, Sacha B. Nelson, Adam Hantman

## Abstract

Uncovering common principles by which diverse modalities of information are processed is a fundamental goal in neuroscience. In mammalian brain, thalamus is the central processing station for inputs from sensory systems, subcortical motor systems, and cortex; a function subserved by over 30 defined nuclei^1,2^. Multiple thalamic nuclei send convergent information to each region of the forebrain, but whether there is a conserved architecture across the set of thalamic pathways projecting to each forebrain area has remained unresolved^3–5^. To uncover organizational principles of thalamic pathways, we produced a near-comprehensive transcriptomic atlas of thalamus. This revealed a common logic for thalamic nuclei serving all major cortical modalities. We found that almost all nuclei belong to one of three major profiles, with a given cortical area getting input from each of these profiles. These profiles lie on a single axis of variance aligned with the mediolateral axis of thalamus, and this axis is strongly enriched in genes encoding receptors and ion channels. We further show that each projection profile exhibits different electrophysiological signatures. Single-cell profiling revealed that rather than forming discrete classes, thalamic neurons lie on a spectrum, with intermediate cells existing between profiles. Thus, in contrast to canonical models of thalamus that suggest it is a switchboard primarily concerned with routing distinct modalities of information to distinct cortical regions, we show that the thalamocortical system is more akin to a molecularly-defined ‘filter bank’ repeatedly applied across modality. Together, we reveal striking covariation in the organization of thalamic pathways serving all input modalities and output targets, establishing a simple and comprehensive thalamic functional architecture.

## MAIN TEXT

To understand the organization of thalamic pathways, we combined anatomical and genetic approaches to produce a near-comprehensive, projection-specific transcriptomic atlas of murine thalamus. Thalamic nuclei were retrogradely labeled from individual forebrain areas, microdissected and cells pooled (8 projection targets, 22 nuclei, 120 samples, Extended Data Tables 1 and 2). Anterograde tracing of inputs to thalamus was used when identification of nuclear boundaries was ambiguous (Fig. 1, A and B, Extended Data Table 1). We then used hierarchical clustering to explore the relationship between the transcriptomes of thalamic nuclei (on the 500 most differentially expressed genes via an ANOVA-like test, see methods, Extended Data Fig. 1B and Supplementary Table 2), and identified five major subdivisions of nuclei across thalamus (Fig. 1C). Anterior dorsal nucleus (AD) and nucleus reuniens (RE) each formed profiles of their own, leaving three major multi-nuclei profiles. These major profiles were not explained by cortical projection target or modality, since the multiple nuclei projecting to motor, somatosensory or visual cortices split across different profiles. For example, central medial (CM), ventral anterior (VA) and ventral lateral (VL) nuclei all project to motor cortex, but are split across the three profiles. By typically receiving input from each of these profiles, each cortical region samples from all three genetically defined pathways. Our nuclear subgrouping also did not obey rules of the prominent core/matric scheme of thalamus, *Calb1*^+^ neurons which define ‘matrix’ nuclei are split by the first branch of our hierarchical clustering (Extended Data Fig. 2)^5–7^. Rather, our three major profiles were best distinguished by anatomy with nuclei of each profile occupying a characteristic position along the mediolateral axis of the thalamus (Fig. 1D). We thus find that the architecture of thalamus is dominated by genetic differences that are organized topographically.

**Fig. 1.**
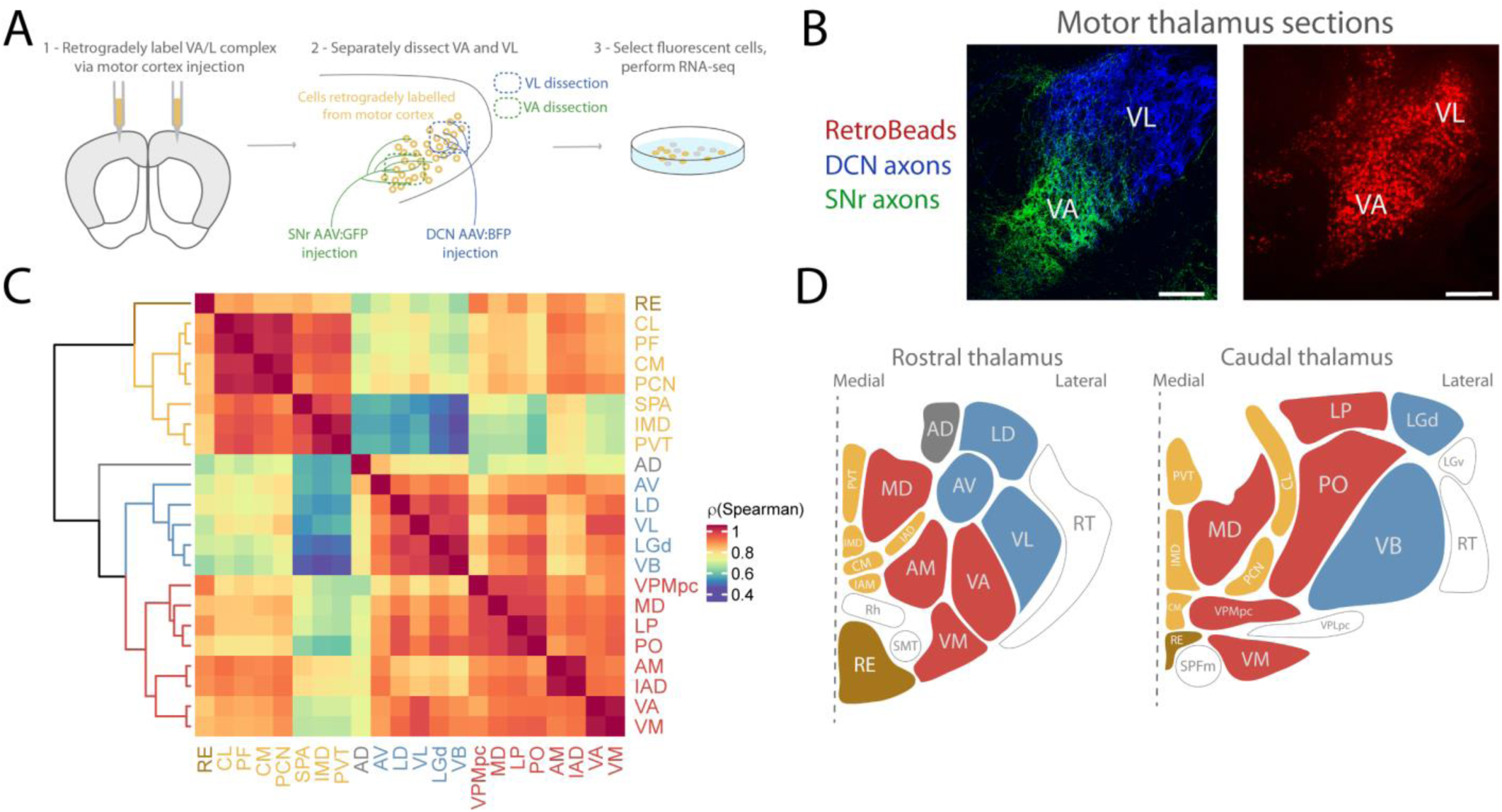
A near-comprehensive transcriptomic atlas allows unbiased clustering of thalamic gene expression profiles. A. Schematic of experimental pipeline to obtain transcriptomic atlas of the thalamus. In this example, motor thalamic neurons were retrogradely labelled from their primary projection target (motor cortex), manually dissected and sorted. Viruses expressing green and blue fluorescent proteins (GFP, BFP), respectively, were injected to the deep cerebellar nuclei (DCN) and substantia nigra pars reticulata (SNr) to label motor nuclear subdivisions (ventral lateral (VL) and ventral anterior (VA), respectively) previously identified^8,9^. B. Example labelling from scheme shown in A. Coronal sections. Scale bars = 200 µm. C. Hierarchical clustering of thalamic nuclei using Spearman’s correlation of top 500 most differentially expressed genes (Extended Data Fig. 1B) across all 22 nuclei. Major clusters defined as the top 5 branches of cluster dendrogram. D. Topographic localization of gene expression profiles in thalamus. Coronal thalamic section schematics with nuclei colored as in Fig. 1C. Unsampled nuclei are left uncolored. (see Fig. 1B).

To understand the pattern of gene expression differences between the thalamic projection profiles, we performed principal component analysis (PCA) on our data. Principal component 1 (PC1, 38% explained variance) separated nuclei into the same major profiles identified via hierarchical clustering (Fig. 2A, Extended Data Fig. 3A). Again, position on this first dimension strongly correlated with mediolateral position, demonstrating topogenetic architecture in thalamus (Fig. 2B, Extended Data Fig. 3B). Based on their relative order on this first component, we named the three major profiles primary, secondary, and tertiary. The progressive difference from primary to tertiary nuclei was also evident in the number of genes differentially expressed between the groups, with the primary and tertiary nuclei being most distinct, and the other two comparisons being less so (Fig. 2C). This primary axis was prominently enriched in genes encoding neurotransmitter receptors, ion channels, and signaling molecules (Fig. 2D and E). Thus, nuclei of the major profiles sit continuously along a single axis of genetic variance which is aligned with the mediolateral spatial axis of thalamus.

**Fig. 2.**
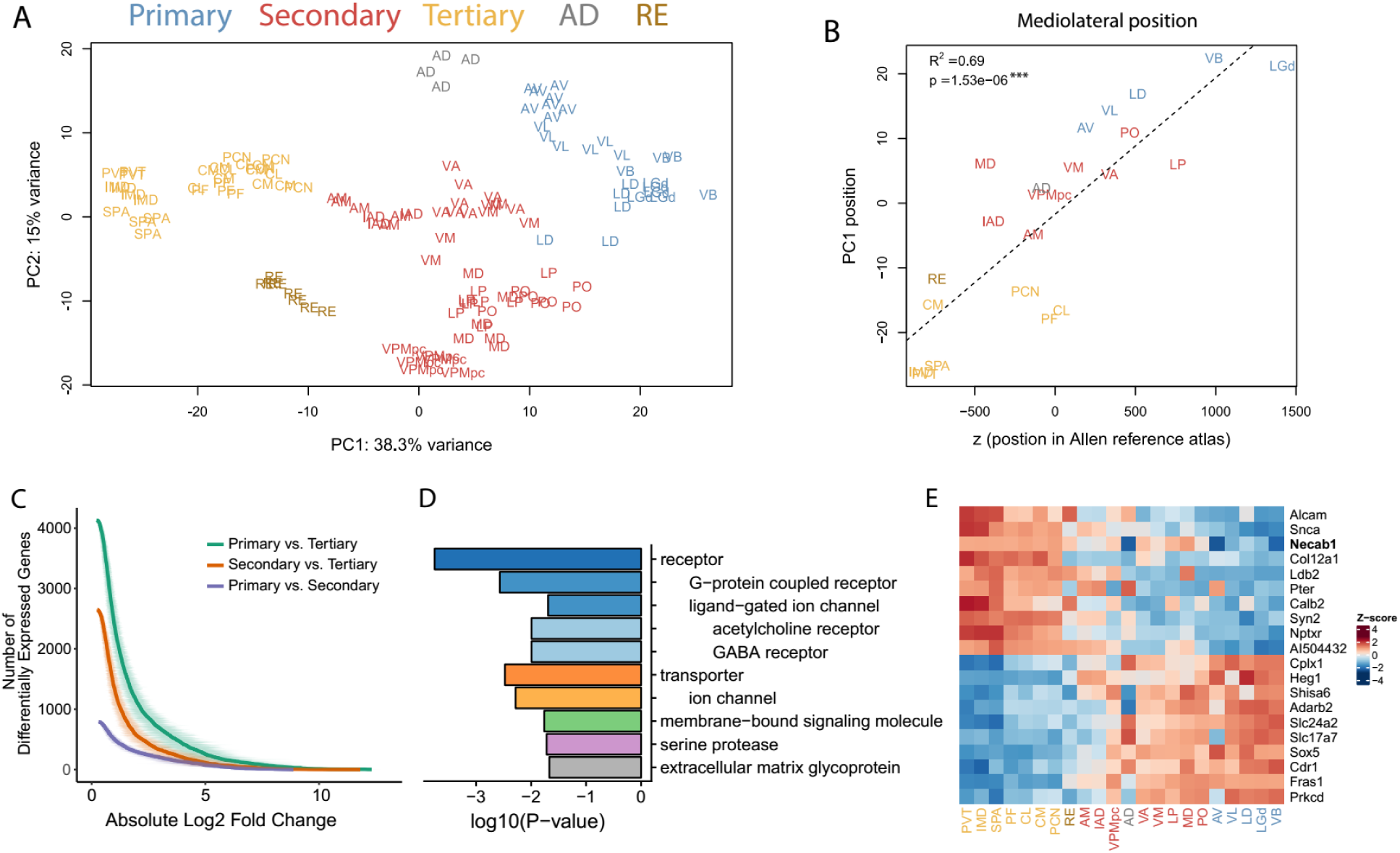
A common topographic axis of variance separates major thalamic profiles. A. PCA showing separation of functional nuclear profiles in the first two principal components. The underlying gene set and color scheme are the same as in 1C. B. PC1 position is highly correlated with mediolateral position of the nuclei. Mediolateral positions are z voxel coordinates of nuclei centers in the Common Coordinate Framework (CCF) in the Allen Mouse Brain Atlas. 1 voxel = 10 µm. C. Primary nuclei are farthest from tertiary nuclei, with secondary nuclei being intermediate. Plot shows the number of differentially expressed genes at each log fold change level (shown as mean ± standard deviation) for the three comparisons. D. Genes relevant to neurotransmission are overrepresented amongst the top 100 genes with the highest absolute PC1 loadings in our dataset. The ten most highly overrepresented PANTHER protein class terms are shown. P-values based on hypergeometric test. Indentation indicates gene subfamily. E. Heatmap of genes with strongest positive and negative loadings on PC1. Nuclei are ordered by their mean position on PC1 of Fig. 2A. Colors represent gene-wise Z-scores.

**Fig. 3.**
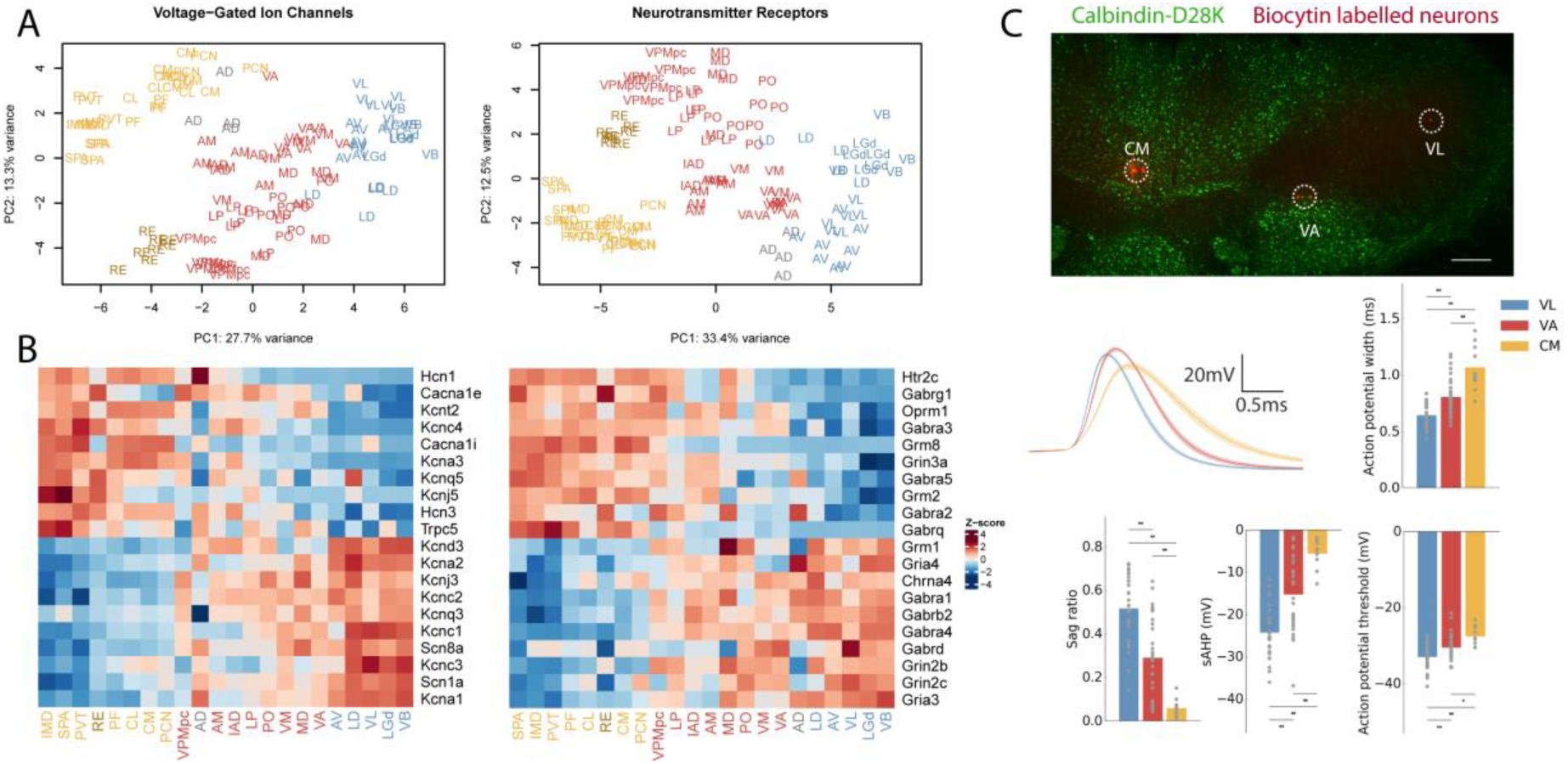
Functionally relevant genes and electrophysiological properties vary systematically across nuclear profiles. A. PCA including only genes encoding voltage-gated ion channels and neurotransmitter/neuromodulator receptors. Colors as in Fig. 1C. B. Heatmap for genes with the highest gene loadings in PC1 from Fig. 3A. Voltage-gated ion-channels on the left and neurotransmitter receptors on the right. Colors represent gene-wise Z-scores. C. Systematic variation of electrophysiological properties across profiles. Whole-cell patch-clamp recordings from VL (primary thalamus), VA (secondary thalamus) and CM (tertiary thalamus). Top row: neurons were labelled with biocytin (red) and localized to individual nuclei with the aid of Calbindin-D28K immunolabelling (green). Scale bar = 100µm. Middle row, left shows average action potential shape for VL, VA and CM neurons (mean±SEM). Remaining panels show comparisons for four physiological measurements across these nuclei (One-way ANOVA with *post-hoc* Tukey HSD test, all comparisons P<0.05). Sample contained 29 VL neurons, 34 VA neurons and 10 CM neurons.

Given the prominent differences in receptor and ion channel expression between thalamic profiles, we asked whether these profiles correspond to functionally distinct classes of neurons. We first performed PCA on the expression profiles of voltage-gated ion channel or neurotransmitter/modulator receptor encoding genes (Fig. 3A, left and right respectively). Analysis with these limited gene sets reproduced the separation of profiles in PC1 (Fig. 3A), confirming that ion channel and receptor profiles are organized along the same axis identified in Fig. 2. Genes linked to high firing rates via faster channel kinetics, such as Kv3 channels (*Kcnc1, Kcnc3*), the *Scn8a* channel, and the *Kcnab3* subunit^12–14^, tended to be progressively elevated toward primary profile nuclei. This raised the possibility that action potential width may progressively narrow from tertiary to primary nuclei (Fig. 3B). Whole-cell recordings from the motor-related nuclei CM, VA and VL (representing the three main nuclear profiles; Fig. 2E) confirmed this prediction (Fig. 3C). Neurons recorded within VL have the narrowest action potential width and those in CM the widest. In addition, many other electrophysiological properties showed a systematic gradient ranging from VL through VA to CM (Fig. 3C, and Extended Data Fig. 4). Prior work has shown substantial differences in electrophysiological properties between different thalamic nuclei, but to date this has not been incorporated into thalamic organizational schemes ^9–11^. Our electrophysiology recordings thus show a systematic variation of neuronal firing properties from primary to tertiary profiles. The strong enrichment of neuromodulatory genes, especially among nuclei of the secondary and tertiary type, suggests further differential modulation of inputs across these profiles.

**Fig. 4.**
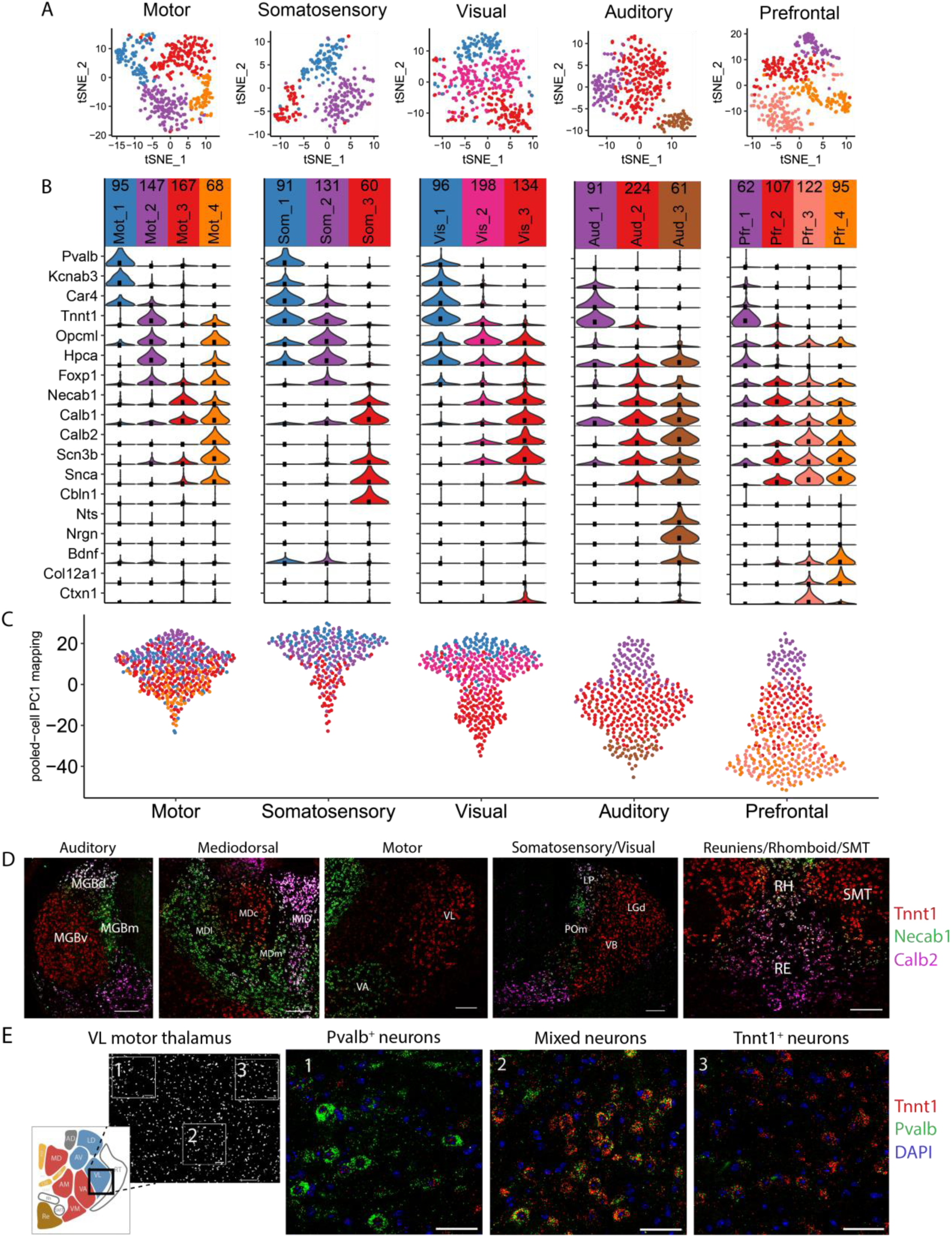
A topographic spectrum of thalamic cell identities within and between thalamic nuclei. A. Overview of single-cell clusters within each projection system visualized via tSNE. Cells colored by cluster identity. B. Violin plots for cluster marker genes (inclusion criteria were Likelihood Ratio Test P-value < 10^−5^ and log2 fold change > 0.5 for each cluster). C. Projection of single-cell data onto pooled-cell PC1 from Fig. 2A. Each dot is a single cell colored as in A. D. Topographic distribution of marker genes within 6 major thalamic modalities. Multi-FISH with probes for *Calb2* (pink), *Tnnt1* (red) and *Necab1* (green). See Extended Data Fig. 7 and 8 for expanded views and quantification. Scale bars = 200 µm. E. Multi-FISH within VL thalamus comparing genes marking clusters in single-cell data. Left panel shows field of view (coronal section, scale bar = 100 µm). Right three panels (scale bars = 50µm) show expansion of the three boxed areas, moving left to right. Middle box shows intermediate cells expressing both single cell cluster markers. (red = *Tnnt1*, green = *Pvalb*, blue = DAPI).

Thus far we analyzed thalamic nuclei by pooling projection neurons from specific anatomical positions. The resulting transcriptional profiles could represent homogenous populations, discrete subtypes, or cells with graded differences. To probe these possibilities, we profiled the transcriptomes of individual neurons from motor, somatosensory, visual, auditory, and prefrontal projection classes (Extended Data Table 3). Analysis of this single-cell RNAseq dataset resulted in multiple clusters for each projection class, and cluster markers included many genes that also distinguished nuclei (Fig. 4B). Single-cell clusters also separated along the first principal component derived from our pooled-cell RNAseq dataset (Fig. 4C, Extended Data Fig. 6). Markers for the single-cell clusters were spatially separated at the single-cell level, but with intermediate cells especially prominent near the anatomical boundaries of thalamic nuclei (Fig. 4D, Extended Data Fig. 7). This is consistent with spatially organized heterogeneity or a gradient-like organization rather than intermingled, discrete populations^15,16^.

Given the strong relationship between PC1 and anatomical nuclei position (Fig. 2C and Extended Data Fig. 3), the presence of single-cell clusters occupying similar PC1 positions (e.g. clusters 1 and 2 of the motor projections neurons, or clusters 1 and 2 of the somatosensory nuclei, Extended Data Fig. 6) suggested that distinct neuron types could also coexist within anatomical boundaries of nuclei. We examined this possibility by performing multi-color fluorescent *in situ* hybridization (multi-FISH) for genes which distinguished amongst the major profiles (*e*.*g. Pvalb* and *Tnnt1*). Taking motor thalamus as an example, *Pvalb* and *Tnnt1* expressing cells were found within the anatomical boundaries of a single thalamic nucleus (VL; Fig. 4E). Some individual VL cells expressed both *Pvalb* and *Tnnt1; Pvalb*-selective, intermediate, and *Tnnt1*-selective cells were distributed along the mediolateral axis (Fig. 4E). Therefore, spatially organized transcriptomic differences can exist even within individual thalamic nuclei.

A common organizing principle of thalamus divides nuclei into discrete core or matrix subtypes that span modalities^1,7^. To date the strongest evidence for cross-modal organization is that genetic differences can be larger within a sensory modality than between sensory modalities^17^(Extended Data Fig. 9). However, previous studies have focused either on select sensory systems and/or have had limited scope with respect to the molecules investigated^18^. Here, using the full transcriptomes of nearly all thalamic pathways, we confirm cross-modal organization but replace the core/matrix dichotomy with a spectrum of profiles that span a single axis of genetic variance. For example, we find that the matrix subtype has substantial diversity, splitting into multiple profiles based upon hierarchical clustering (Extended Data Fig. 2). The axis of variance is dominated by genes that directly shape neuronal properties, leading to conserved, systematic variation of function. This pattern of variation is not only imposed on sensory thalamocortical systems but also diversifies motor, limbic, and cognitive thalamocortical systems. Understanding how the pattern of intra-thalamic molecular variability intersects with input modalities and behavioral relevance will be an important challenge for future work^19–23^.

Our single-cell transcriptomics indicated that a given cortical area samples across a spectrum of thalamic profiles. Some projection neurons exhibited features of more than one profile, suggesting gradient-like transitions between cell types. Using multi-FISH, we mapped these intermediate cell types both to the boundaries between nuclei and within nuclei. Intermediate cells might exhibit hybrid input-output transforms, morphologies, and/or functions.

Given that neurons spanning the full range of profiles provide input to nearly all cortical areas, our data suggests that thalamus acts as a “filter bank” providing each recipient forebrain area with a broad complement of differentially filtered inputs (Extended Data Fig. 10).

## Materials and Methods

### EXPERIMENTAL METHODS

#### Animal care

Experimental procedures were approved by the Institutional Animal Care and Use Committee (IACUC) at the Janelia Research Campus. Mice were housed on a 12 hour light/dark cycle, with *ad libitum* food and water. The majority of mice were 8-12 weeks old (Supplementary Tables 1 and 3).

#### Acquisition of samples

Cells were fluorescently labelled to enable manual dissection. This was done through retrograde labelling via either viral or tracer injection into the major projection field of the nucleus of interest. For viral injections, rAAV2-retro expressing cre-dependent GFP or TdTomato under the CAG promoter were injected, with volumes of 50-100nL at 3 depths (see Extended Data Table 2) ^24^. Minimum survival time was 3 weeks post-injection. Viruses were prepared by Janelia Virus Services. Non-viral retrograde tracer labelling used the lipophilic tracer DiI (2.5mg/ml in DMSO, injecting volumes of 50-200nL per site, from Molecular probes) or Lumafluor red retrobeads (diluted 3x in PBS, 50-200nL per site). Anterograde labelling of inputs to thalamus was also used in a small number of cases (see Extended Data Tables 1 and 2).

We referred to the Paxinos and Franklin mouse brain atlas to guide our dissections^25^. For the majority of regions of thalamus, retrograde tracers labeled populations corresponding to identified thalamic nuclei (Extended Data Tables 1 and 2 for targeting details). However, the caudal intralaminar nuclei (parafascicular complex) were less clearly delineated. This likely reflects additional heterogeneity within this complex beyond that shown in atlases^26^.

At no stage were experimenters blinded to sample identity.

#### Manual cell sorting and RNAseq

##### Sorted pooled-cell RNAseq

Fluorescent cells were collected and sequenced as previously described^15,27^. Briefly, animals were deeply anaesthetized with isoflurane and euthanized. Coronal slices (200-300 μm) were cut and placed for 1 hour at room temperature with pronases and neural activity blockers in artificial cerebrospinal fluid (ACSF). Relevant regions were then microdissected, and the tissue dissociated. The resulting cell suspensions were diluted with filtered ACSF and washed at least 3 times by transferring them to clean dishes. This process produces negligible contamination with non-fluorescent tissue (Extended Data Fig. 1A)^27^. After the final wash, samples were aspirated in a small volume (3 μl) and lysed in 47μl XB lysis buffer using the Picopure kit (KIT0204, ThermoFisher) in a 200μl PCR tube (Axygen), incubated for 30 min at 40 °C on a thermal cycler and stored at -80 °C.

Library preparation and sequencing was performed by the Janelia Quantitative Genomics core. RNA was isolated from each sample using the PicoPure RNA isolation kit (Life technologies) and on-column RNAase-free DNase I treatment (Qiagen). 1µL ERCC RNA spike-in mix (Life technologies) was added to each sample. Amplification was then performed using the Ovation RNA-seq v2 kit (NuGEN), yielding 4-8 µg of cDNA. The Ovation rapid DR multiplexing kit (NuGEN) was used to make libraries for sequencing, which were sequenced on a HiSeq 2500 (Illumina).

##### Sorted single-cell RNAseq

Retrogradely labeled cells were isolated as described above, and collected into 8-well strips containing 3 µL Smart-seq2 lysis buffer, flash-frozen on dry ice, and stored at -80°C until further use^28^.

Upon thawing, cells were re-digested with Proteinase K and barcoded RT primers were added. cDNA synthesis was done using the Maxima H Minus RT kit (Thermo Fisher) and E5V6NEXT template switch oligo, followed by heat inactivation reverse transcriptase. PCR amplification using the HiFi PCR kit (Kapa Biosystems) and SINGV6 primer was performed with a modified thermocycling protocol (98°C for 3 min, 20 cycles of 98°C for 20s, 64°C for 15s, 72°C for 4 min, final extension at 72°C for 5 min). Samples were then pooled across strips, purified with Ampure XP beads (Beckman Coulter), washed twice with 70% ethanol and eluted in water. These pooled strips were then combined to create the plate-level cDNA pool for tagmentation, and concentration was determined using Qubit High-Sensitivity DNA kit (Thermo Fisher).

Tagmentation and library preparation using 600 pg cDNA from each plate of cells was then performed with a modified Nextera XT (Illumina) protocol, but using the P5NEXTPT5 primer and tagmentation time extended to 15 minutes(*30*). The libraries were then purified following the Nextera XT protocol (at 0.6X dilution) and quantified by qPCR using Kapa Library Quantification (Kapa Biosystems). 6-10 plates were run on a NextSeq 550 flow cell. Read 1 contained the cell barcode and unique molecular identifier (UMI). Read 2 was a cDNA fragment from the 3’ end of the transcript.

#### Multi-FISH

C57Bl/6J mice (∼8 weeks old) were anesthetized with isoflurane then fixed via transcardial perfusion with PBS followed by 4% paraformaldehyde in PBS, pH 7.4. Brains were post-fixed at 4 °C overnight, washed 3 times with PBS, and cryoprotected in a sucrose series of 10%, 20% then 30% in PBS at 4 °C. All solutions were prepared RNase-free. Brains were sectioned (14µm) on a Leica CM3050S cryostat, mounted onto Fisher SuperFrost Plus slides, and stored at -80 °C. Multi-FISH was performed using the RNAscope Multiplex Fluorescent Assay platform from ACDBio, following the manufacturer’s protocol. The probes used were: *Calb2* (ref 313641-C3), *Necab1* (ref 428541 and 428541-C2), and *Tnnt1* (ref 466911-C2). Fluorescent dyes were DAPI, Alexa Fluor 488, Atto 550 and Atto 647. Images were acquired using a Zeiss LSM 880 confocal microscope. Images were acquired with an air 20x (0.8NA) objective unless otherwise specified.

#### Electrophysiology

Acute brain slices were prepared from p20-25 mice. Animals were deeply anesthetized with ketamine/xylazine/acepromizine and transcardially perfused with ice-cold oxygenated cutting solution containing (in mM): 74 NaCl, 3 KCl, 1 NaH2PO4, 25 NaHCO3, 6 MgCl2, 0.5 CaCl2, 5 Sodium Ascorbate, 75 Sucrose, 10 Glucose. 300 µm coronal slices containing the thalamus were cut on a vibratome (Leica), and then recovered for 15 min at 33 °C and for 15 min at room temperature in oxygenated cutting solution followed by at least another 1 hour at room temperature in oxygenated ACSF containing (in mM): 126 NaCl, 3 KCl, 1 NaH2PO4, 25 NaHCO3, 2 MgCl2, 2 CaCl2, 10 Glucose. During recordings, slices were perfused with oxygenated 34-35 °C ASCF with 35 mM d,l-2-amino-5-phosphonovaleric acid (APV), 20 mM 6,7-dinitroquinoxaline-2,3-dione (DNQX) to block glutamatergic synaptic transmission and 50 mM picrotoxin to block GABAergic synaptic transmission. Target neurons in CM, VA and VL were identified based on their distance to the mammillothalamic tract and nuclear borders were confirmed with calbindin immunostaining *post hoc*. Whole-cell recording pipettes (6 – 8 MΩ) were filled with internal solution containing (in mM): 100 K-gluconate, 20 KCl, 10 HEPES, 4 Mg-ATP, 0.3 Na-GTP, 10 Na-phosphocreatine, and 0.1% biocytin. Current-clamp recordings were obtained with Multiclamp 700B amplifiers (Molecular Devices) digitized at 10 kHz using IGOR Pro (WaveMetrics). Resting membrane potentials were adjusted to -65 mV and steady state series resistance was compensated. Neurons with series resistance > 30 MΩ or membrane potentials that changed by > 3 mV were excluded.

Custom IGOR scripts were used to analyze the data. For each neuron, threshold, amplitude, afterhyperpolarization and half width at half height of the 16th-19th action potentials in trials with 20 to 40 Hz firing rate were averaged.

#### Immunohistochemistry

After recordings, slices were fixed with 4% paraformaldehyde and 2.5% sucrose in 5x phosphate-buffered saline (PBS) at 4 °C for 2-10 days. After washing with PBS, slices were blocked in PBS with 0.3% Triton and 3% BSA at 4 °C overnight and then incubated in PBS with 0.3% Triton and 3% BSA and rabbit anti-calbindin D-28k (Swant, 1:1000) at 4 °C overnight. After washing, they were incubated in PBS with 0.3% Triton, 3% BSA and 5% goat serum with fluorescent protein conjugated goat anti-rabbit IgG (Invitrogen, 1:1000) and streptavidin (Invitrogen, 1:1000) at 4 °C overnight to label calbindin-expressing neurons and biocytin-filled neurons.

### ANALYSIS METHODS

#### Pooled-cell RNAseq analysis

##### Data processing and quality control

After removing Illumina adapter sequences using cutadapt, reads were mapped to the mouse reference genome (mm10) using STAR with ‘ENCODE settings’ for RNAseq^29^. Mean mapping rate was 82.29% with a standard deviation of 2.25%. Unique unambiguous exon-mapping reads were summarized at the gene level using Gencode version M13.

Contamination with common astrocytic, oligodendrocytic, erythrocytic and microglial transcripts was low, consistent with a lack of substantial contamination by non-fluorescent cells (Extended Data Fig. 1A). To ensure the specificity of our dissections and to control for potential batch effects, we collected several nuclei through multiple independent labelling approaches, and showed that these samples cluster in a highly similar manner (Extended Data Fig. 1C).

##### Differential gene expression

Differential expression was assessed using the Bioconductor package *edgeR*^30^. Low counts were removed by requiring a Transcripts per million (TPM) > 5 in at least 3 samples. This yielded a list of approximately 17,000 expressed genes. Counts were then fitted to a negative binomial generalized linear model, where each factor level represents a different thalamic nucleus, and a Likelihood Ratio Test was used to assess differential expression between groups. P-values were adjusted for multiple tests using the Benjamini-Hochberg method. Genes with false-discovery rate < 0.05 were considered differentially expressed. For selecting the most differentially expressed genes between any thalamic nuclei, we used an ANOVA-like test (ANODEV test for generalized linear models, as described in edgeR User manual 3.2.6), testing for differences between any of the 22 nuclei, and used the 500 genes with the highest P-value. To avoid bias due to differences in sample number when comparing numbers of differentially expressed genes between different profiles in Fig. 2C, the groups were subsampled (with replacement) to the size of the smallest group. Bootstrapped log2 fold changes were obtained over 100 iterations.

For visualization, clustering and machine learning of gene expression data, we used variance-stabilized counts produced by the variance-stabilizing transformation in the *DESeq2* R package)^31,32^.

For assessing the role of modality vs. hierarchical class on distinguishing thalamic nuclei, we used elastic-net regularized logistic regression classifiers. Models were trained with different numbers of randomly selected genes as features over 100 iterations. To avoid bias due to variable group size, groups were subsampled to the size of the smallest group. Model tuning was performed using the *glmnet* and *caret* packages in R, and accuracy of the best model was assessed using 5-fold cross-validation.

##### Unsupervised clustering and functional enrichments

Hierarchical clustering was performed using 1 - Spearman’s correlation as a distance metric and complete linkage for agglomeration. Groups were defined by splitting the tree at the level of 5 branches. We termed these profiles, not clusters, as we do not mean to imply discreteness between the classifications. PCA was done using the singular value decomposition based *prcomp* function in R. For functional enrichment of differentially expressed genes, we used the PANTHER Protein Class Ontology (http://data.pantherdb.org/PANTHER13/ontology/Protein_Class_13.0), which is a consolidated version of molecular function gene ontology. Over-representation in the top 100 genes with the highest PC1 loadings was assessed via hypergeometric test.

For defining voltage-gated ion channels and neurotransmitter receptors, we downloaded the IUPHAR/BPS database (http://www.guidetopharmacology.org/DATA/targets_and_families.csv). Voltage-gated ion channels were the genes defined in the database, while for neurotransmitter receptors we included ionotropic and metabotropic receptors for glutamate, GABA, glycine, acetylcholine, 5-HT, dopamine, trace amine, histamine, and opioids.

#### Single-cell RNAseq analysis

##### Data processing and quality control

Single-cell RNAseq data was trimmed for adapters using cutadapt and aligned to the mouse genome (mm10) using STAR. To demultiplex cells, collapse UMIs and produce gene-wise counts for each cell, we used a modified version of the *Drop-seq_tools-1*.*12* pipeline. Briefly, read 1 was tagged based on the cell barcode and UMI, and this information was added to read 2 by merging back the reads after mapping, followed by gene-wise tagging of reads that map onto exons and summarization of digital counts.

Single cells were required to have more than 20,000 UMIs and more than 2,500 genes detected per cell, which yielded a total of 1,971 cells (Extended Data Fig. 5A). Of these, 22 cells were found to be significantly contaminated with oligodendroglial and vascular cell transcripts (Extended Data Fig. 5C), leaving 1,949 cells for all downstream analyses. Genes were considered expressed if their expression was detected in more than 10 cells.

Our single-cell sequencing was not comprehensive, and with improved sequencing approaches further genetic subdivisions may be identified. Single-cell and pooled-cell dissections were not precisely matched, for example motor-projecting midline nuclei were not dissected for single-cell RNAseq. However, pooled-cell and single-cell RNAseq are in close agreement (Extended Data Fig. 4B), indicating that our results are robust to collection method.

##### Single-cell clustering and marker genes

Single-cell clusters were defined using the *Seurat* R package (version 2.0) ^33,34^. Data were log transformed and scaled. For identifying variable genes, genes were divided into 20 bins based on average expression, and genes that were more than 1 standard deviation away from average dispersion within a bin were used for downstream analysis. Single-cell clustering was performed separately for each projection system using shared nearest neighbor (SNN) clustering and limiting the analysis to the top 10 principal components for distance calculation. Clusters were defined by the Louvain algorithm, and clustering resolution set to 0.6. Clusters of cells were visualized using t-distributed stochastic neighbor embedding (tSNE) using the top 10 principal components as input and perplexity set to 30. Marker genes for each cluster were required to be expressed in at least 80% of the cells in the cluster, to have a P-value <10^-5 (Likelihood Ratio Test), a log2 fold change > 0.5. Projection of single-cell data onto pooled-cell principal components was obtained by multiplying (dot product) log-transformed and scaled single-cell data by the pooled-cell principal component loadings.

## End Notes

### Funding

This project was funded as a small project team (ThalamoSeq) by HHMI at the Janelia Research Campus, following a pilot project in the Dudman/Hantman labs. SN and CL were also supported by grants from NINDS (NS079419) and NIMH (MH105949). AS is funded via the Janelia Visiting Scientists Program.

### Contributions

JP: contributed to all aspects of this project. AS: Analyzed and collected data, planned project and wrote the paper. EH: Planned project and collected data. CL: Collected and analyzed electrophysiology data. LW: Collected data. BS: Collected data. WK: Supervised project, AL: Collected data and developed methods JD: Supervised project and wrote the paper SN: Supervised project. AH: Initiated and supervised project, wrote the paper.

### Competing interests

Authors declare no competing interests.

### Data and materials availability

All transcriptomic data used will be made publicly available via the Gene Expression Omnibus.

### Further acknowledgements

We thank Karel Svoboda, Albert Lee and Amy Chuong for critical input throughout the project. We thank Matthew Phillips, Kirandeep Ghataorhe, Brett Mensh and Yves Weissenberger for comments on the manuscript. We thank Vilas Menon, Damian Kao and Mark Cembrowski for help with single-cell RNAseq analysis. We thank Kim Ritola and the Janelia Virology and histology cores for production of viruses and histology. We thank Daniel Morozoff, Yajie Llang, Justin Little, Amy Chuong and Na Ji for surgical protocols and assistance identifying nuclei for dissection, and the Janelia vivarium services for animal care and surgeries.

## Extended data figures

**Extended Data Fig. 1.**
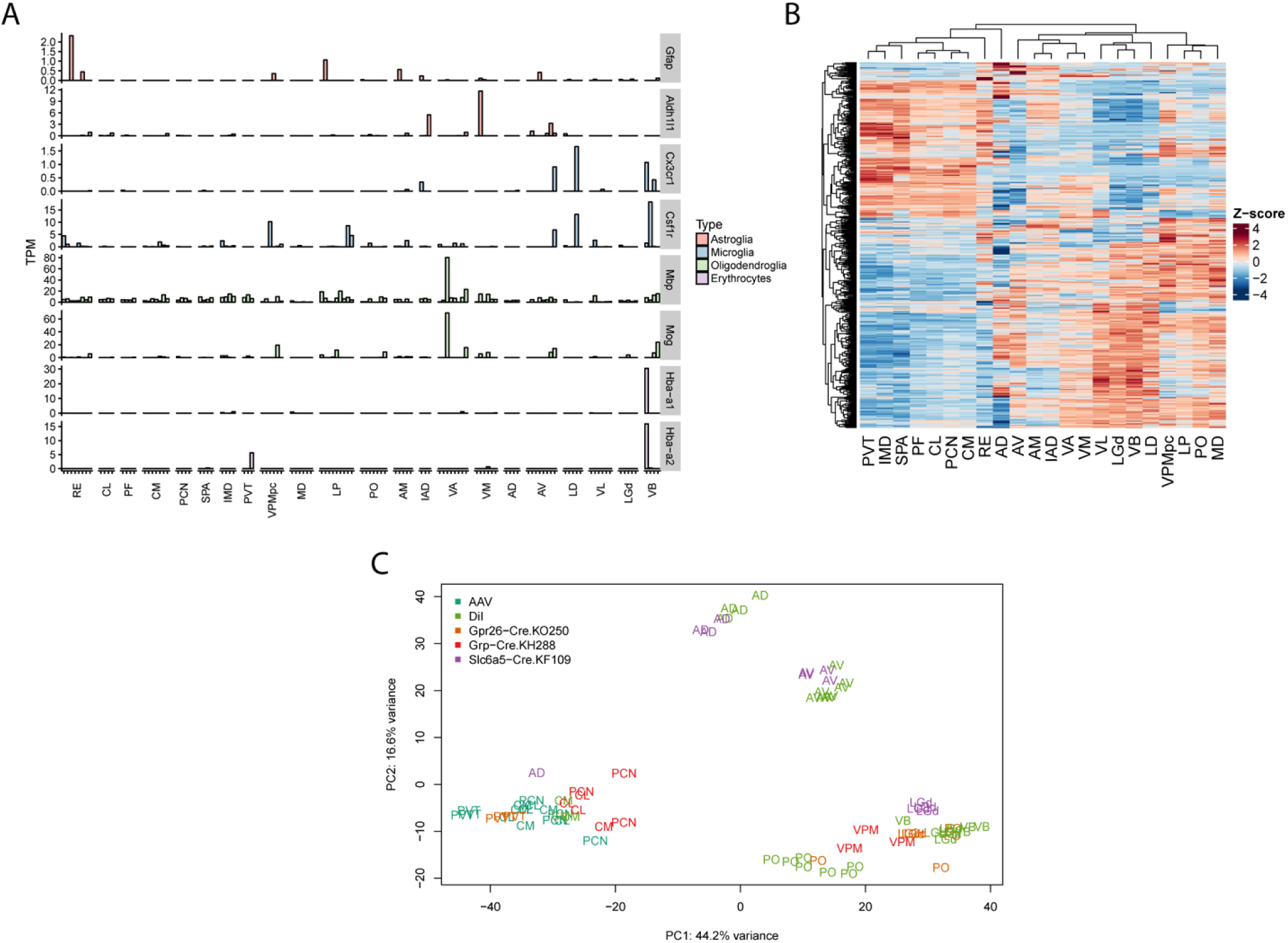
Pooled-cell RNAseq quality control and additional analyses. A) Markers of non-neuronal sample contamination are low across our dataset. Expression (TPM) in pooled-cell samples shown for 8 genes marking astrocytes, microglia, oligodendrocytes and erythrocytes. Only a small number of samples showed expression of contamination markers. B) Heatmap of the top 500 differentially expressed genes. Rows and columns are ordered by hierarchical clustering with Euclidean distance metric. Colors represent gene-wise Z-scores. C) Samples of the same nucleus obtained via different labelling methods cluster similarly. Principal components analysis of those samples, for which multiple collection methods were used (i.e. GENSAT lines in addition to retrograde labeling) using the top 500 genes with highest variance. Samples are colored by collection approach or transgenic line used.

**Extended Data Fig. 2.**
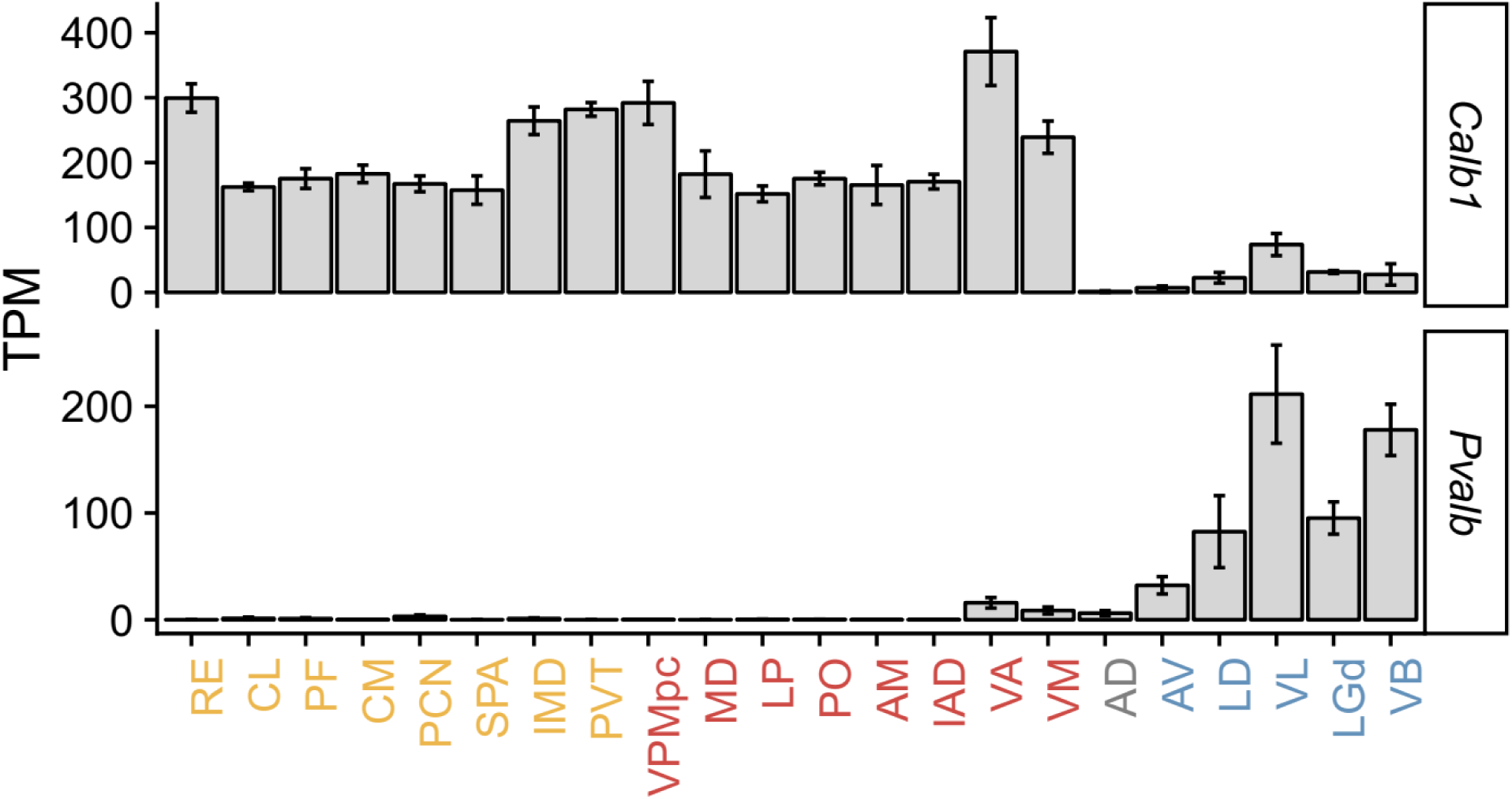
Existing thalamic marker genes do not cleanly mark nuclear profiles. Expression of *Calb1* and *Pvalb* (mean ± SEM) for each nucleus, with nuclei on the x-axis colored by their profile from Fig. 1. The core/matrix organizational theory proposes that thalamus is divided into two discrete groups, expressing calbindin or parvalbumin. However, the first major branch splits the secondary and tertiary groups, both of which are marked by *Calb1* and would thus both be considered ‘matrix’ nuclei in this theory. Thus existing markers for thalamic nuclei subgroups do not adequately reflect thalamic organizational structure.

**Extended Data Fig. 3.**
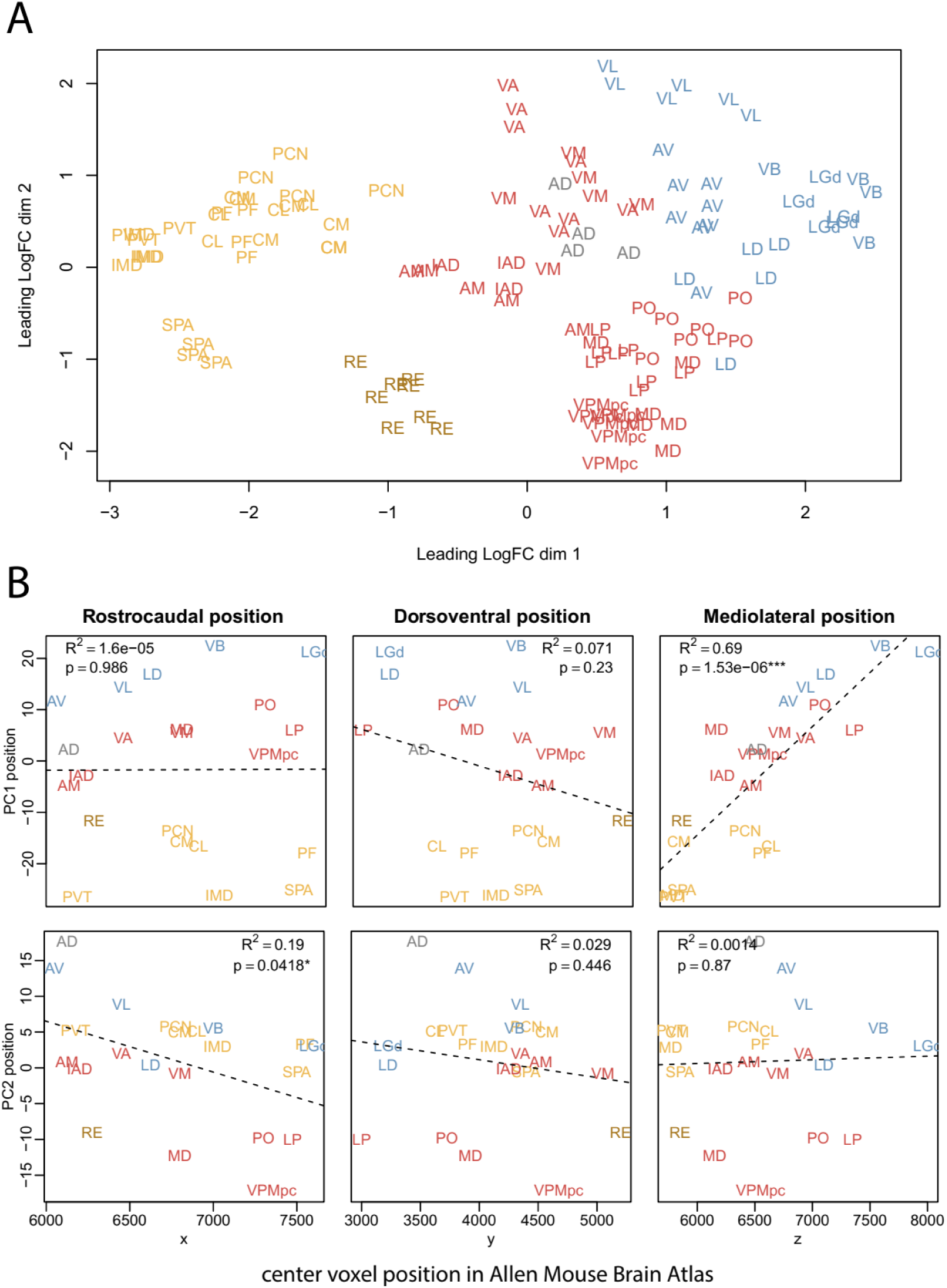
Additional analysis of topogenetic axis. A) Multidimensional scaling using an alternative distance metric also identifies the same leading axis of variance. Distance was defined as the quadratic mean of the log2 fold changes of the top 500 differentially expressed genes between any two samples (meaning that the gene set used for the distance comparison varies between each sample comparison made). B) Relationship of PC1 and 2 with topographical position of nuclei. Rostrocaudal, dorsoventral, and mediolateral positions are the x, y, and z voxel coordinates, respectively, in the Allen Mouse Brain Atlas. 1 voxel = 10µm.

**Extended Data Fig. 4.**
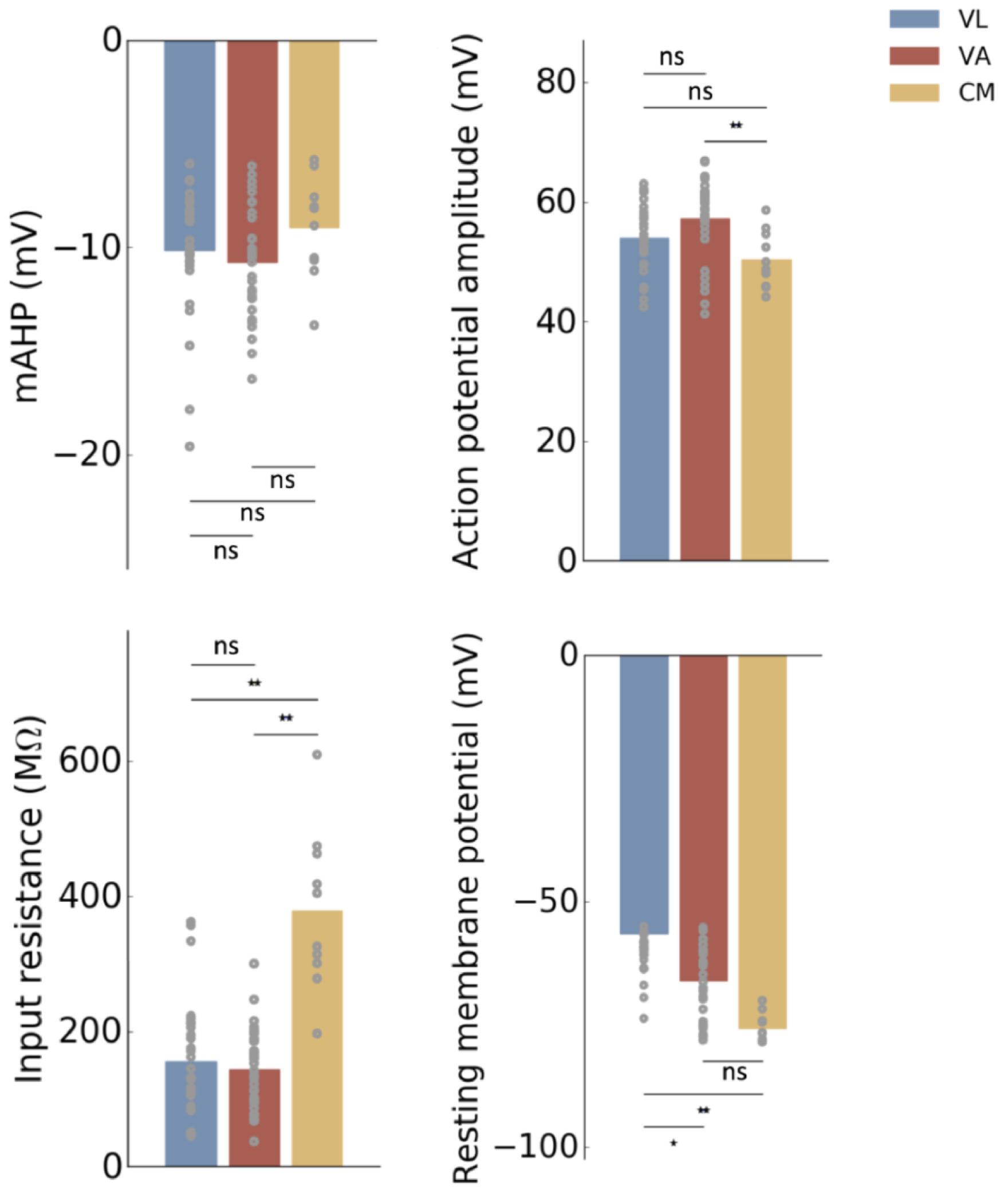
Additional electrophysiological properties between thalamic nuclear profiles. All statistical tests and experimental details are the same as in figure 3C.

**Extended Data Fig. 5.**
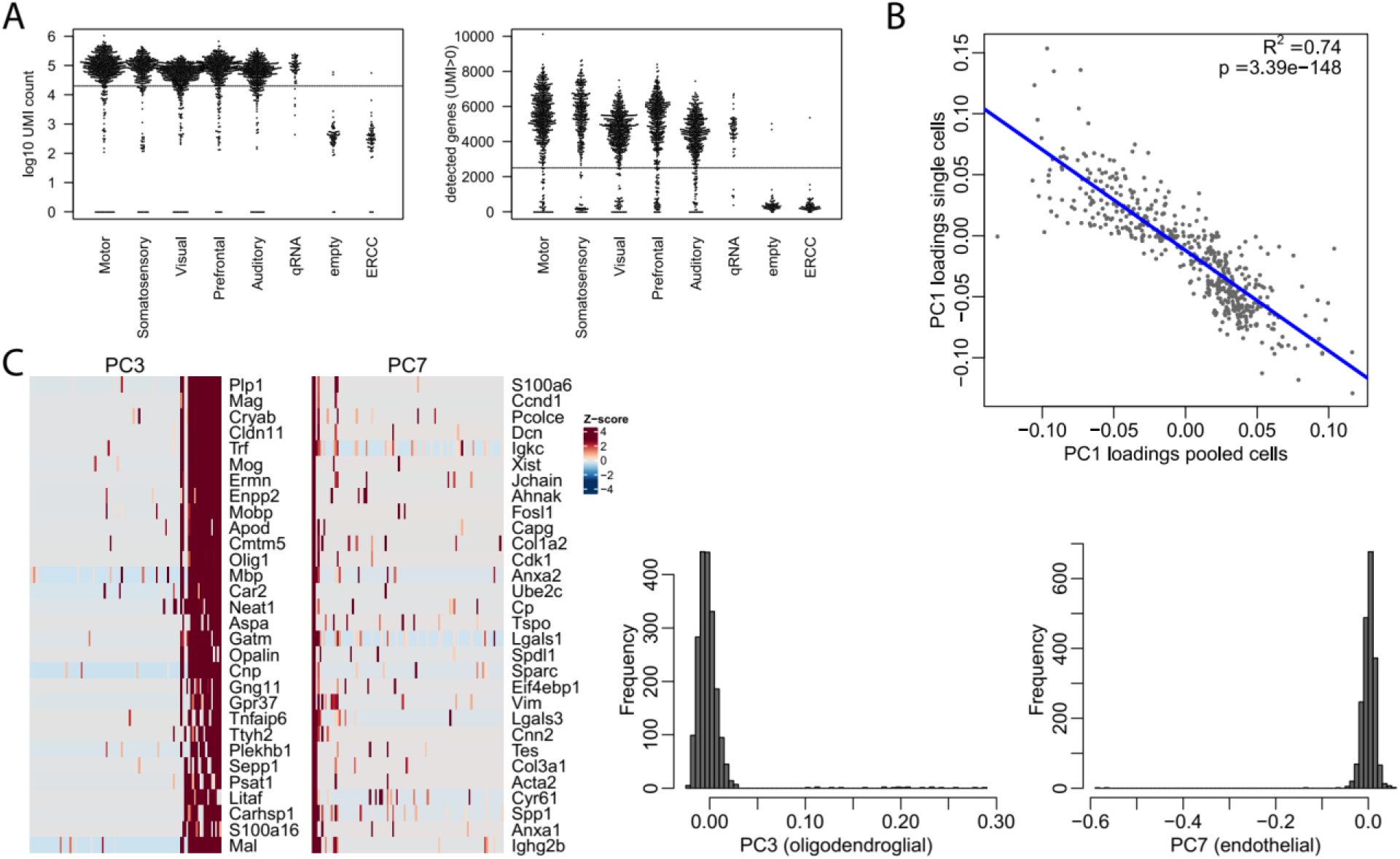
Quality control for single-cell RNAseq data. A) UMI count (left) and gene detection rate (right) for all collected single cells. Cutoffs for downstream use are indicated by dashed lines. B) PC1 loadings for the most differentially expressed genes between nuclei (gene set as in Fig.1) are highly similar in pooled-cell and single-cell RNAseq data. C) PCA on the single-cell RNAseq data revealed that principal components 3 and 7 represented non-neuronal contamination from oligodendrocytes and endothelial cells. Cells with PC3 position >0.05 and PC7 position <0.1 were removed.

**Extended Data Fig. 6.**
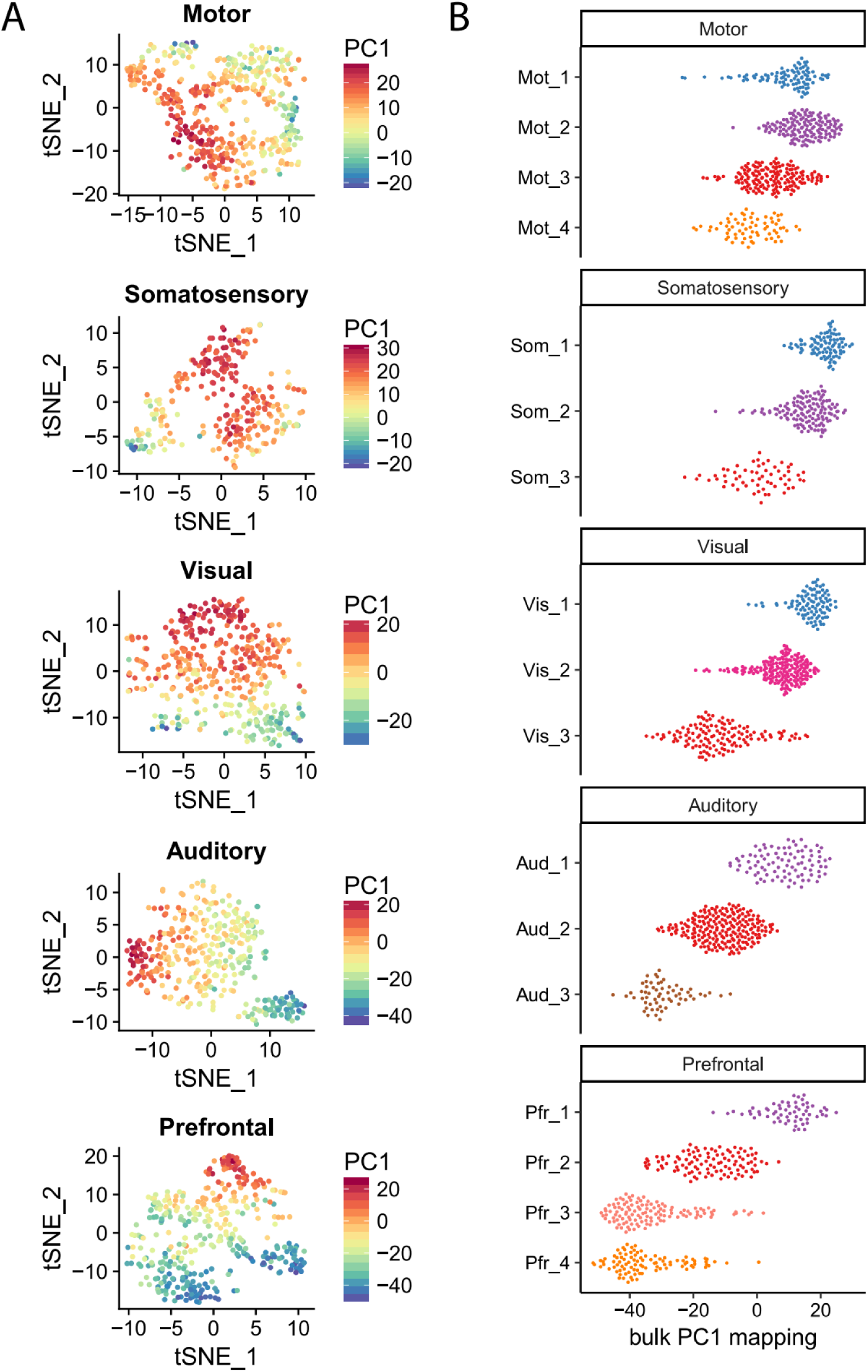
Projection of single cells onto PC1 of pooled-cell RNAseq data. A) tSNE plots for each projection type with cells from single-cell RNAseq colored by their projection onto pooled-cell PC1. B) Positions of single cells projected onto pooled-cell PC1 from Fig. 2, plotted separately for each single-cell cluster.

**Extended Data Fig. 7.**
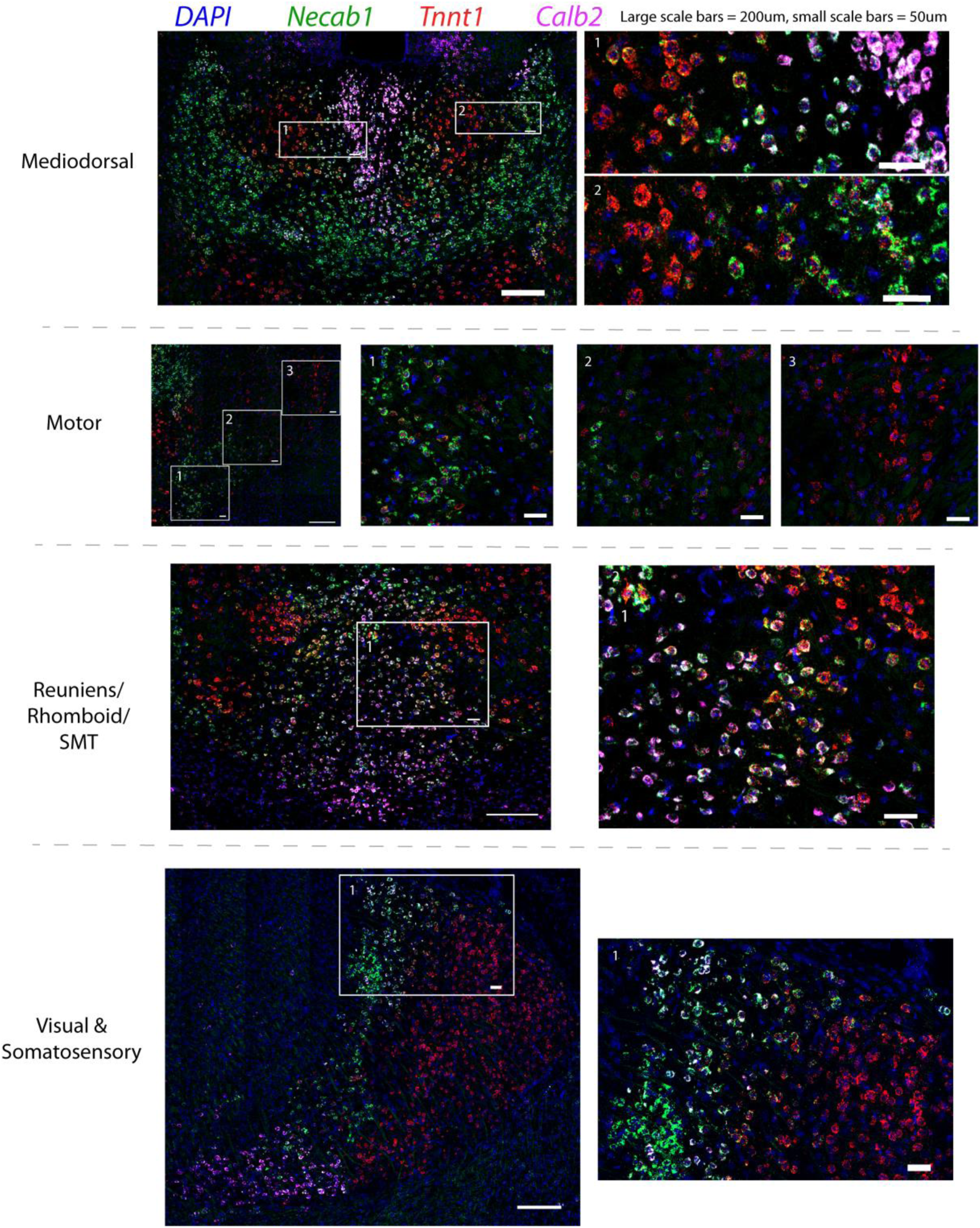
Multi-FISH show cells with mixed expression of profile marker genes. Expanded views of example regions showing intermediate cells expressing combinations of *Tnnt1, Necab1* and *Calb2*, which are preferentially expressed in primary, secondary and tertiary nuclear profiles respectively.

**Extended Data Fig. 8.**
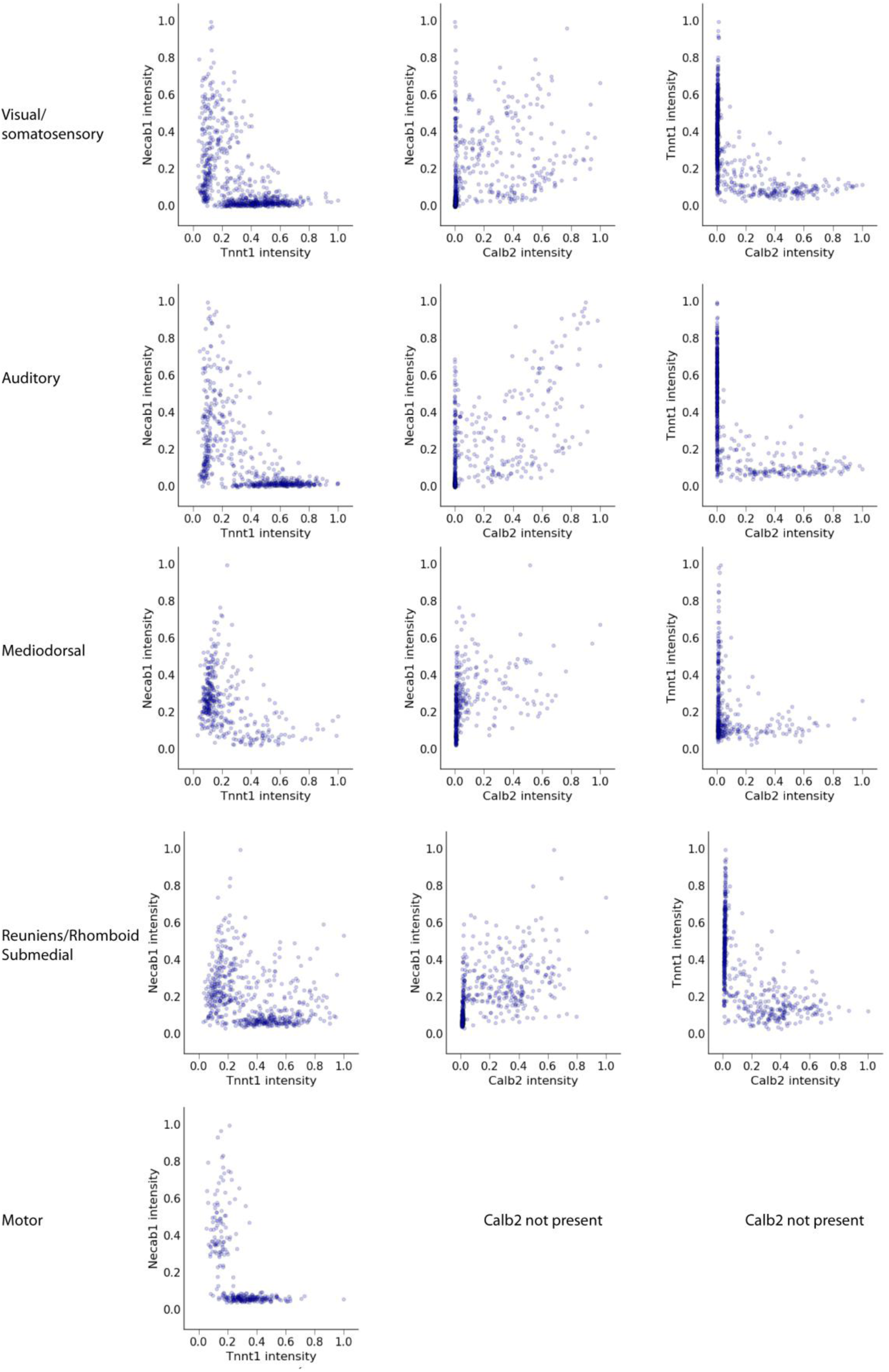
Quantification of multi-FISH images shows intermediate cells. Quantification of multi-FISH gene expression images. Regions of interest (ROIs) were drawn in ImageJ. Intensity was normalized first to the ROI size, then divided by the maximum for that channel. Only cells that express at least one of the marker genes were included.

**Extended Data Fig. 9.**
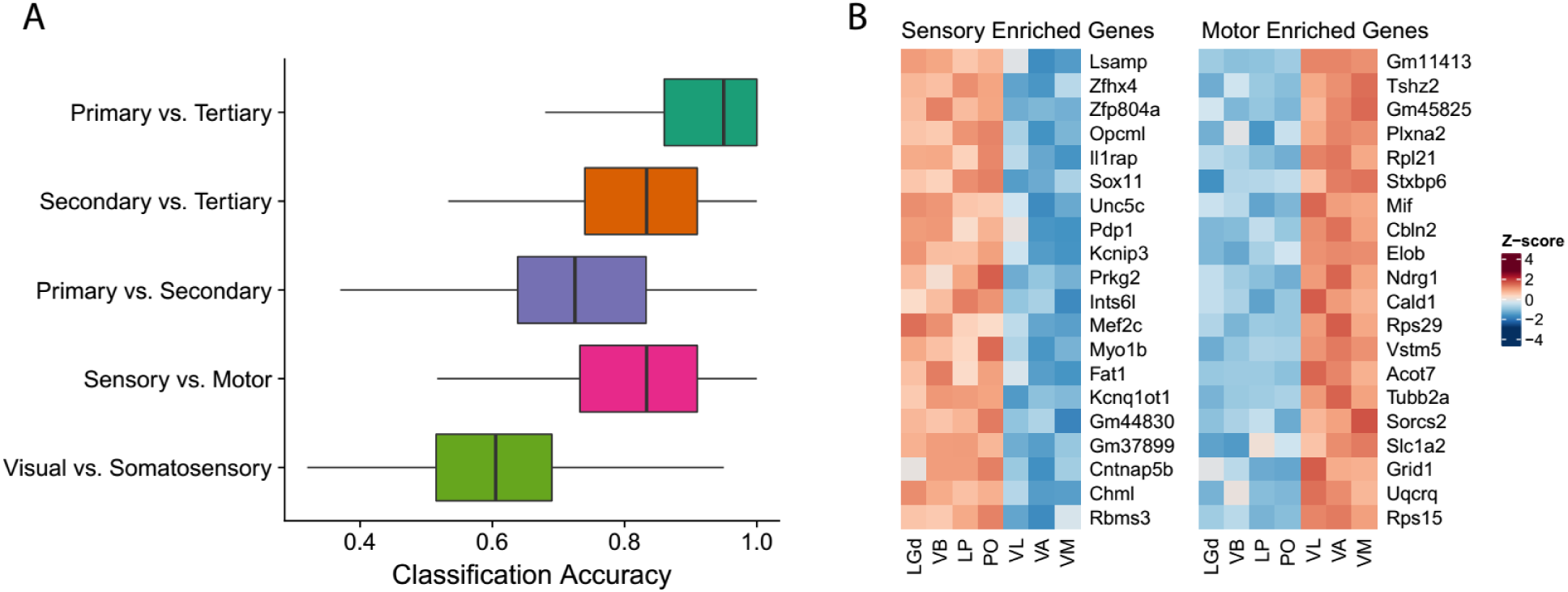
Differential gene expression between sensory and motor thalamic nuclei. A) Classification accuracy for distinguishing Primary, Secondary, and Tertiary type nuclei, as well as for distinguishing motor (VL,VA,VM) vs. sensory (LGd,LP,VB,PO) and visual (LGd,LP) vs. somatosensory (VB,PO) nuclei. Classifiers were obtained using elastic-net logistic regression on 20 random genes over 100 iterations. To prevent bias due to sample size difference, groups were subsampled to the size of the smallest group (n=9) at each iteration. Accuracy was assessed using 5-fold cross-validation. A value of 1 corresponds to perfect classification, while 0.5 corresponds to chance level performance. B) Genes that best distinguish motor from sensory nuclei (LGd,VB,LP,PO vs. VL,VA,VM). Plotted are the top 20 genes with false discovery rate < 10^−3^ (Likelihood Ratio Test), fold change > 2, and ordered by highest signal-to-noise ratio (mean log fold change between vs. within group).

**Extended Data Fig. 10.**
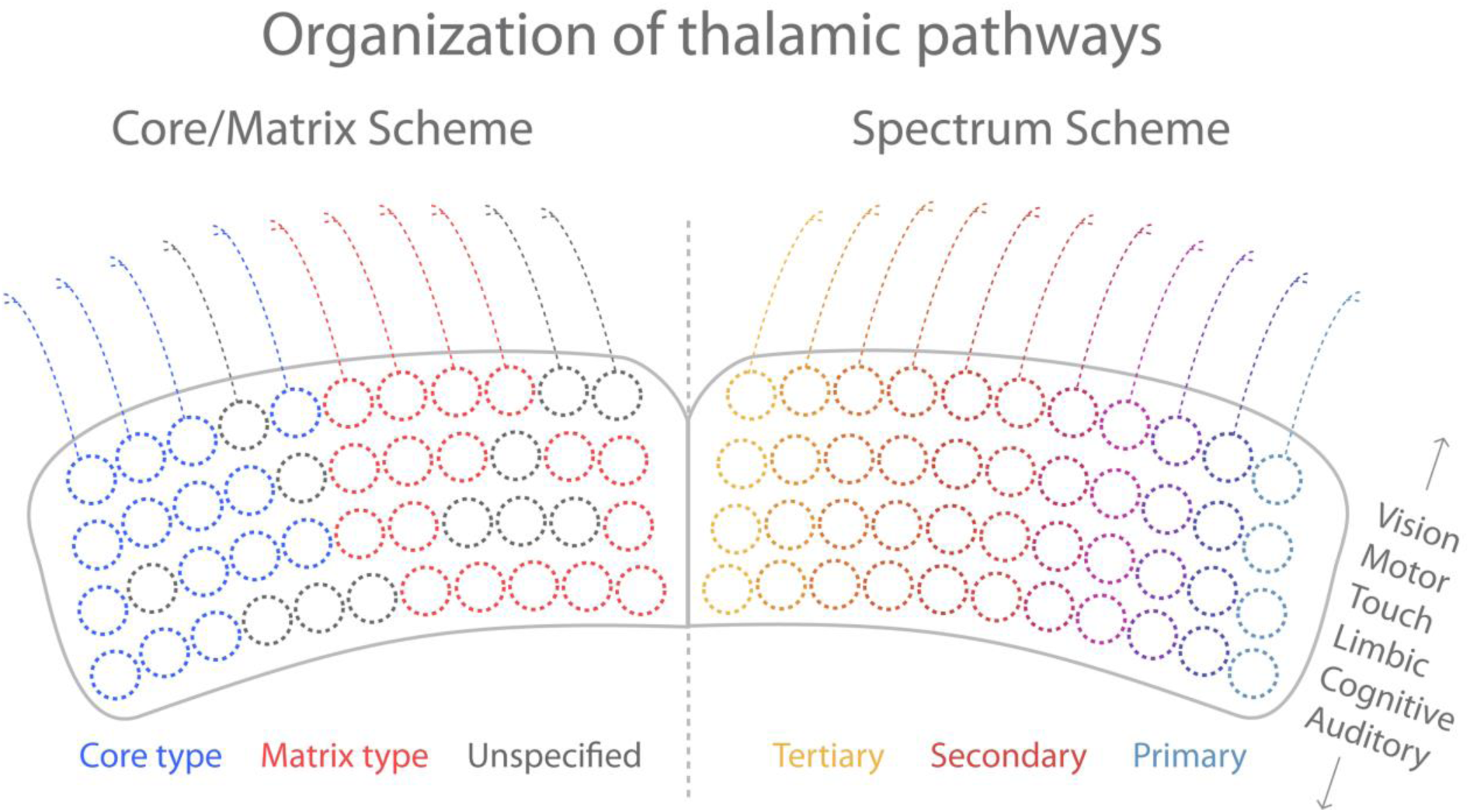
A spectrum of thalamic pathways conserved across modalities. Left: The core/matrix scheme divides thalamus into two discrete cell types based upon their expression of parvalbumin (Core nuclei) and calbindin (Matrix nuclei). However, many nuclei do not express either marker (for example, the anterior nuclei), and calbindin expressing nuclei can have markedly different anatomical and functional characteristics (for example, the rostral intralaminar nuclei and PO both are calbindin^+^). Right: The spectrum scheme identifies three major profiles of nuclei, but places them on a single spectrum. This spectrum is aligned with the mediolateral axis of thalamus, is based upon the entire transcriptome, and links projection type directly to functional properties. Thalamus therefore consists of a set of filter banks for each modality.

## Extended Data Tables

**Extended Data Table 1.**
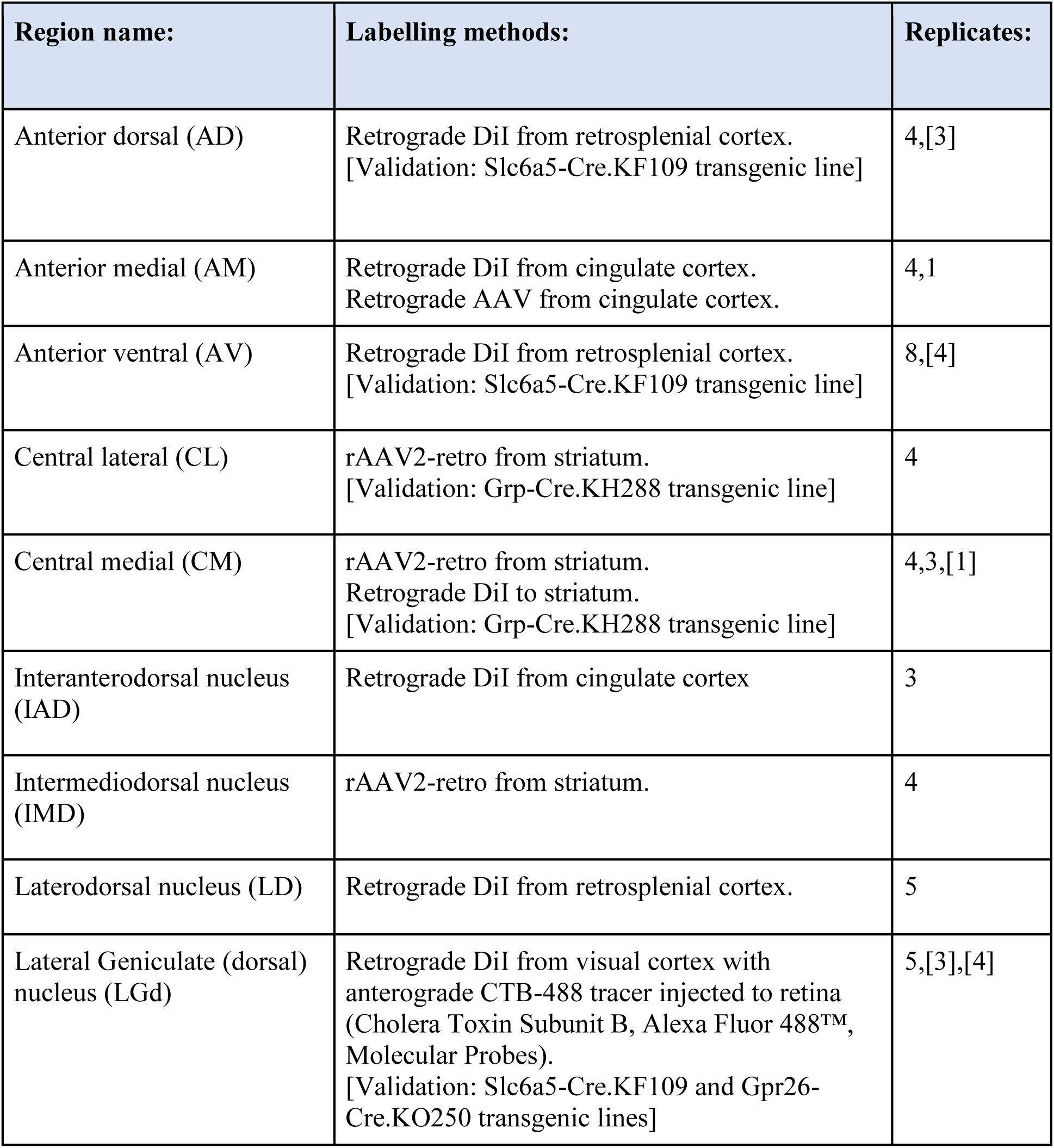
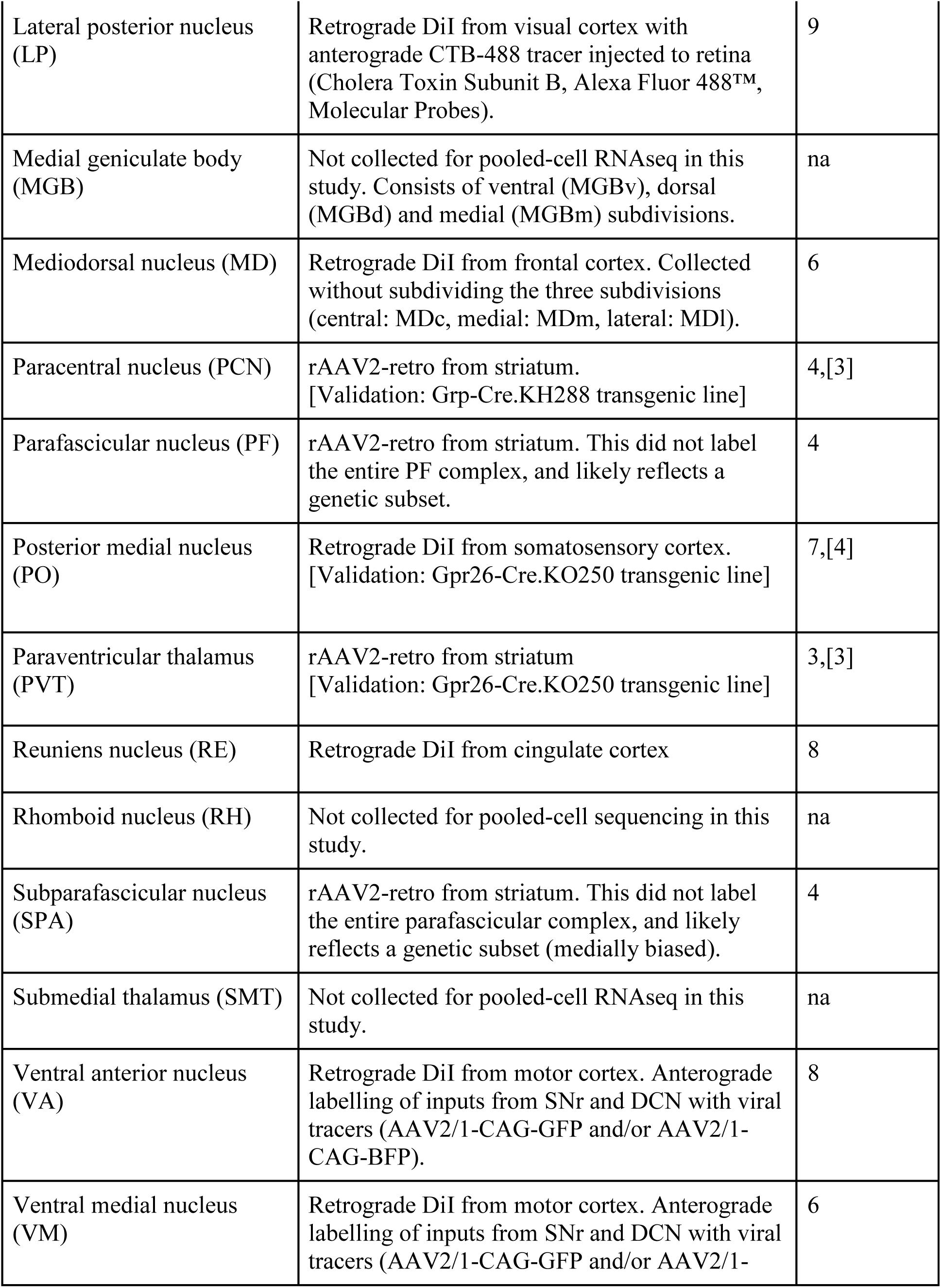
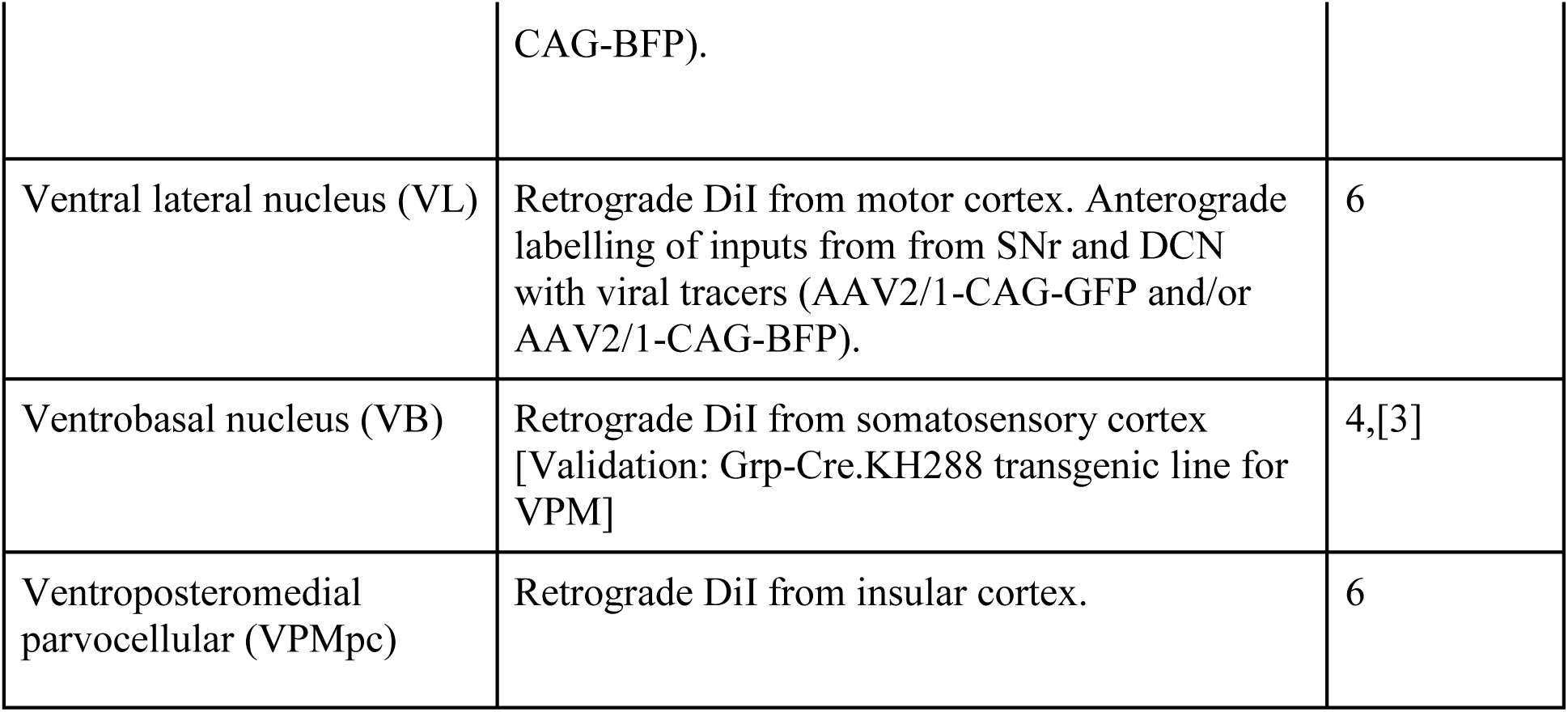
Sample collection approaches (pooled-cell RNAseq) [Validation] refers to samples in Extended Data Fig. 1C. For transgenic lines used, see supplementary Extended Data Table 4.

**Extended Data Table 2.**
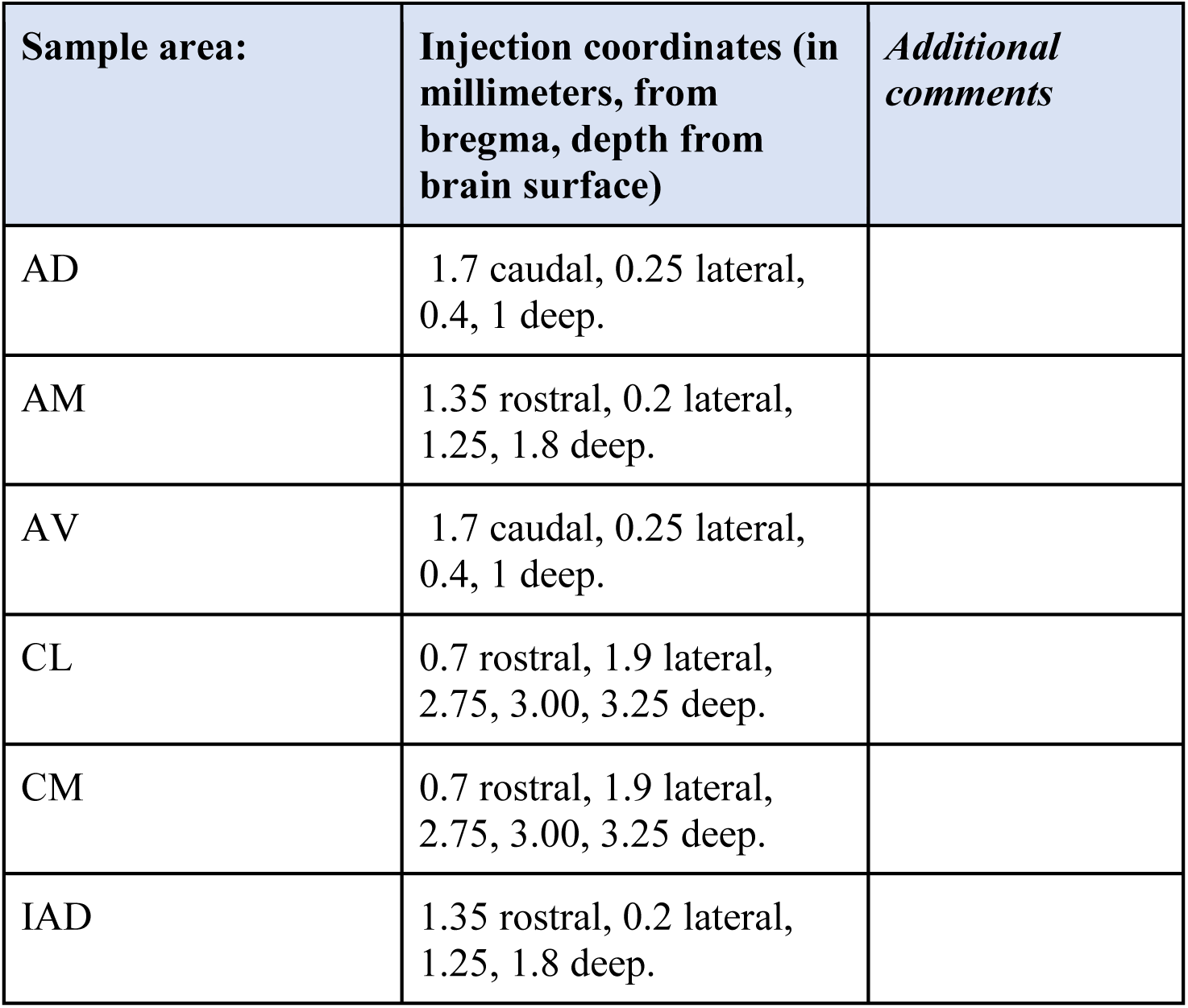
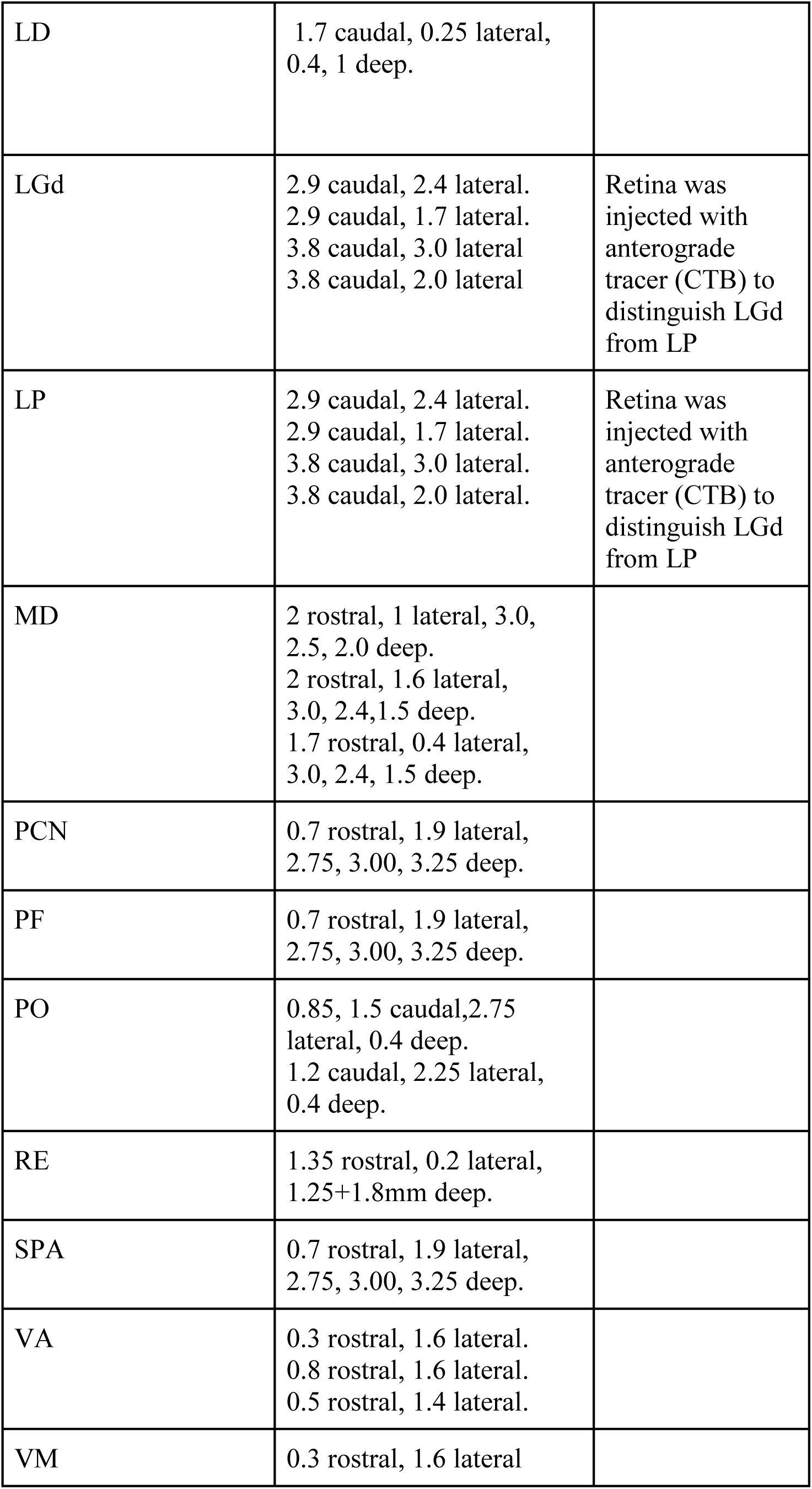
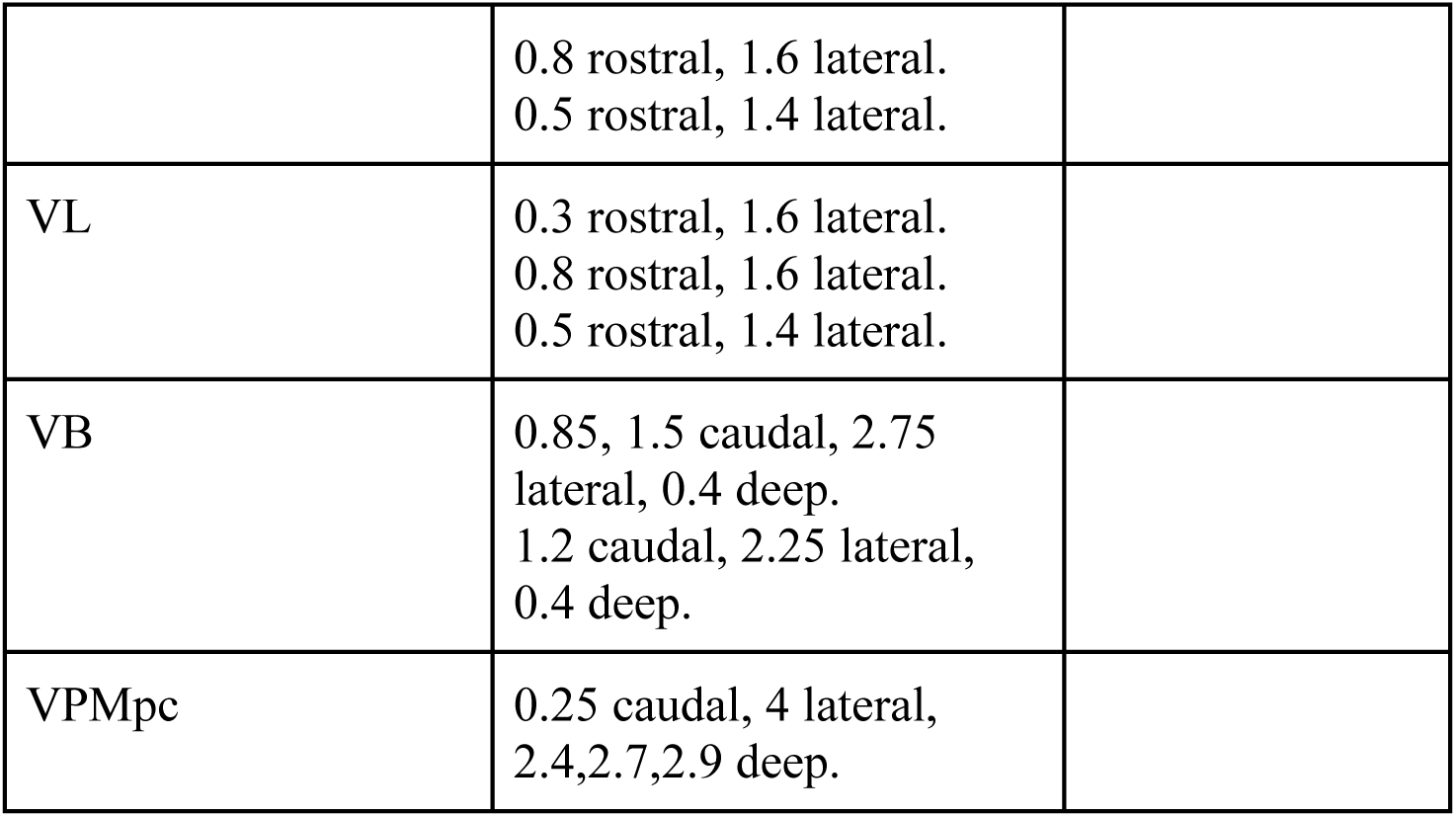
Coordinates for retrograde labelling/trace injections (pooled-cell RNAseq) All depths relative to brain surface. If depth not stated, injections were made at 300µm and 600µm deep.

**Extended Data Table 3.** Injection coordinates for retrograde labelling (single-cell RNAseq)

| **Sample area:** | **Injection coordinates (in millimeters, from bregma, depth from brain surface)** |
| --- | --- |
| Auditory Thalamus | 3.5 caudal, 4.5 lateral, 2.25 deep.<br>3.16 caudal, 4.5 lateral, 2.0 deep. |
| Mediodorsal (Prefrontal) Thalamus | 2 rostral, 1 lateral, 3.0, 2.5,2.0 deep.<br>2 rostral, 1.6 lateral, 3.0,2.4,1.5 deep.<br>1.7 rostral, 0.4 lateral, 3.0, 2.4, 1.5 deep. |
| Motor Thalamus | 0.3 rostral, 1.6 lateral.<br>0.8 rostral, 1.6 lateral.<br>0.5 rostral, 1.4 lateral. |
| Somatosensory Thalamus | 0.85 and 1.5 caudal, 2.75 lateral, 0.4 deep.<br>1.2 caudal, 2.25 lateral, 0.4 deep. |
| Visual thalamus | 2.9 caudal, 2.4 lateral.<br>2.9 caudal, 1.7 lateral.<br>3.8 caudal, 3.0 lateral.<br>3.8 caudal, 2.0 lateral. |

**Extended Data Table 4.** Transgenic mice used in this study.

| Transgenic line: | Areas collected |
| --- | --- |
| Slc6a5-Cre.KF109 (GENSAT) | Anterior dorsal, Anterior ventral, Dorsal lateral geniculate (LGd). |
| Gpr26-Cre.KO250. (GENSAT) | Paraventricular nucleus (PVT), Posterior medial (PO) |
| Grp-Cre.KH288 (GENSAT) | Rostral intralaminar nuclei (Central lateral, paracentral), ventral posteromedial nucleus (VPM). |

### Captions for Supplementary Tables

**Supplementary Table 1** (separate file) – Pooled-cell RNAseq metadata and differential gene expression.

**Supplementary Table 2** (separate file) – Pooled-cell RNAseq principal component analysis and PANTHER protein class enrichment.

**Supplementary Table 3** (separate file) – Single-cell RNAseq metadata and cluster marker genes.

